# Structural characterization and AlphaFold modeling of human T cell receptor recognition of NRAS cancer neoantigens

**DOI:** 10.1101/2024.05.21.595215

**Authors:** Daichao Wu, Rui Yin, Guodong Chen, Helder V. Ribeiro-Filho, Melyssa Cheung, Paul F. Robbins, Roy A. Mariuzza, Brian G. Pierce

## Abstract

T cell receptors (TCRs) that recognize cancer neoantigens are important for anti-cancer immune responses and immunotherapy. Understanding the structural basis of TCR recognition of neoantigens provides insights into their exquisite specificity and can enable design of optimized TCRs. We determined crystal structures of a human TCR in complex with NRAS Q61K and Q61R neoantigen peptides and HLA-A1 MHC, revealing the molecular underpinnings for dual recognition and specificity versus wild-type NRAS peptide. We then used multiple versions of AlphaFold to model the corresponding complex structures, given the challenge of immune recognition for such methods. Interestingly, one implementation of AlphaFold2 (TCRmodel2) was able to generate accurate models of the complexes, while AlphaFold3 also showed strong performance, although success was lower for other complexes. This study provides insights into TCR recognition of a shared cancer neoantigen, as well as the utility and practical considerations for using AlphaFold to model TCR–peptide–MHC complexes.

## Introduction

Adoptive cell therapy (ACT) with tumor-specific T cells has been shown to promote durable regression of diverse cancers, including metastatic melanoma, cervix, breast, bile duct, and colon cancers (1–6). The therapeutic effect of these *ex vivo*-expanded tumor-infiltrating lymphocytes (TILs) is mediated mainly by cytotoxic CD8^+^ T cells (7). The principal target of tumor-specific T cells are neoantigens arising from somatic mutations in self-antigens during malignant transformation (6, 8). Of special interest for ACT are neoantigens derived from oncogenes such as *KRAS* and *TP53* that bear driver mutations because these mutations are tumor-specific and essential for cancer cell fitness and proliferation (9).

In a pioneering clinical study of ACT, a patient with metastatic colorectal cancer was effectively treated with four CD8^+^ T cell clones specific for a neoepitope arising from the KRAS^G12D^ driver mutation presented by HLA-C*08:02 (2). All metastases that retained HLA-C*08:02 expression regressed. In another study, a patient with chemorefractory breast cancer was treated with a T cell clone that targeted the R175H driver mutation in the p53 oncogene (10). The patient experienced 55% tumor regression for 6 months.

RAS proteins are binary switches that play a causal role in many human cancers. They comprise a small GTPase that alternates between an inactive GDP-bound and an active GTP-bound state which regulates cell survival, growth, and differentiation (11). The three main RAS isoforms, KRAS, NRAS, and HRAS, share identical 86-residue N-terminal sequences. This region contains three mutational hotspots, at positions G12, G13, and Q61. Mutation of G12 or G13 prevents the arginine finger of GTPase-activating proteins (GAPs) from entering the GTPase active site and promoting hydrolysis (11). Mutation of Q61, which is part of the GTP hydrolysis mechanism, destroys both intrinsic and GAP-mediated GTP hydrolysis, thereby rendering RAS proteins persistently active.

KRAS is the most highly mutated RAS isoform across cancers, accounting for ∼85% of all RAS mutations with especially high frequencies in pancreatic and colorectal cancers (12, 13). However, NRAS mutations dominate in melanomas, with mutations at position 61 appearing in ∼20% of patients (14). The two most frequent mutations are Q61K and Q61R (15). Mutant NRAS melanomas have more aggressive clinical features and poorer outcomes compared to non-mutant NRAS melanomas (16–18). The immunogenicity of NRAS mutations in melanoma patients was demonstrated by the detection of T cell responses against the NRAS^Q61K^ neoantigen (15). Several T cell receptors (TCRs) were isolated from TILs of these patients that recognize a robustly presented neoepitope corresponding to residues 55–64 of NRAS^Q61K^ that contains the glutamine-to-lysine mutation at position 61 (ILDTAG**K**EEY; mutant amino acid in bold). One of the TCRs also recognizes the NRAS^Q61R^ neoepitope, which contains a glutamine-to-arginine mutation at position 61 (ILDTAG**R**EEY), although peptide titrations indicated that a higher concentration of the NRAS^Q61R^ peptide was required. The TCRs are restricted by the prevalent MHC class I allele HLA-A*01:01 (15). These oligoclonal TCRs, transduced into a patient’s peripheral blood lymphocytes for ACT, may prove effective in eliminating tumors expressing HLA-A*01:01 and the NRAS^Q61K^ or NRAS^Q61R^ mutation. Other recent work has identified a TCR mimic antibody that targets NRAS^Q61R^ and HLA-A*01:01, underscoring the interest in that neoantigen as an immunotherapeutic target (19).

McShan et al. (20) recently determined the solution structure of the NRAS^Q61K^–HLA-A*01:01 complex by nuclear magnetic resonance (NMR) and used molecular dynamics (MD) simulations to probe the conformational dynamics of the MHC-bound NRAS^Q61K^ peptide. This study showed that the side chains of T58, K61, and E63 form an exposed molecular surface, whereas those of I55, L56, and Y64 are buried in the peptide-binding groove of HLA-A*01:01. However, understanding how TCRs discriminate between wild-type and mutated NRAS requires knowing the structure of TCR–peptide–MHC (TCR–pMHC) complexes (21). Here we report crystal structures of a TCR (N17.1.2) from a melanoma patient that recognizes both NRAS^Q61K^ and NRAS^Q61R^ neoepitopes (15) in complex with NRAS^Q61K^–HLA-A*01 and NRAS^Q61R^–HLA-A*01, as well as the structure of the unbound TCR.

The deep learning method AlphaFold (22) has shown impressive performance in predictive modeling (23) and has been adapted and tested for modeling TCR–pMHC complex structures (24, 25), which as noted recently can potentially be utilized for large-scale T cell specificity prediction (26). Given that immune recognition (including TCR–pMHC recognition) is generally not as accurately modeled as other protein complexes by AlphaFold (27), and that others have noted concerns about AlphaFold’s accuracy and utility in some scenarios (28), we tested the capability of AlphaFold to model TCR N17.1.2 in complex with its neoantigen targets, and also tested it for predictive modeling of complexes for other TCRs known to bind NRAS^Q61K^–HLA-A*01. Interestingly, AlphaFold generated accurate models of both N17.1.2 complexes, but accuracy depended on AlphaFold implementation and the number of models produced, whereas it did not generate high scoring models for the other complexes modeled. These findings establish the structural basis for T cell recognition of NRAS^Q61^ neoantigens, and provide valuable insights into the accuracy and limitations of AlphaFold in a challenging predictive modeling scenario.

## Results

### TCR N17.1.2 Discriminates between Mutant and Wild-type NRAS Peptides

TCR N17.1.2 was isolated by screening TILs from melanoma patients for reactivity towards the mutated NRAS^Q61K^ neoantigen (15). This HLA-A*01:01-restricted TCR recognizes the NRAS^Q61K^ and NRAS^Q61R^ neoepitopes using gene segments TRDV1 and TRAJ27-1 for the α chain and TRBV27 and TRBJ2-5 for the β chain. We used surface plasmon resonance (SPR) to measure the affinity of TCR N17.1.2 for HLA-A1 loaded with wild-type or mutant NRAS peptides (**Fig. 1**). Recombinant TCR and pMHC proteins were expressed by *in vitro* folding from bacterial inclusion bodies. Biotinylated wild-type NRAS–HLA-A1, NRAS^Q61K^–HLA-A1, or NRAS^Q61R^–HLA-A1 was directionally coupled to a streptavidin-coated biosensor surface and different concentrations of N17.1.2 were flowed sequentially over the immobilized pMHC ligand. TCR N17.1.2 bound mutant NRAS neoantigens with dissociation constants (*K*_D_s) of 1.2 μM for NRAS^Q61K^–HLA-A1 and 3.4 μM for NRAS^Q61R^–HLA-A1 (**Fig. 1*A*, *B***). These affinities are well within the range of TCRs specific for microbial antigens (*K*_D_ = 1–50 μM) (29) and are comparable to those of TCRs recognizing other cancer neoantigens (21). No apparent interaction between TCR N17.1.2 and wild-type NRAS–HLA-A1 was detected, even after injecting high concentrations (up to 200 μM) of TCR (**Fig. 1*C***). The exquisite specificity of N17.1.2 for NRAS^Q61K^ and NRAS^Q61R^ compared to wild-type NRAS, as measured by SPR, is consistent with functional assays showing that T cells transduced with these TCRs can be activated by APCs pulsed with sub-nanomolar concentrations of mutant NRAS^Q61K^ and NRAS^Q61R^ peptides, but do not respond to wild-type NRAS peptide, even at >1000-fold higher concentrations (15).

**Fig. 1.**
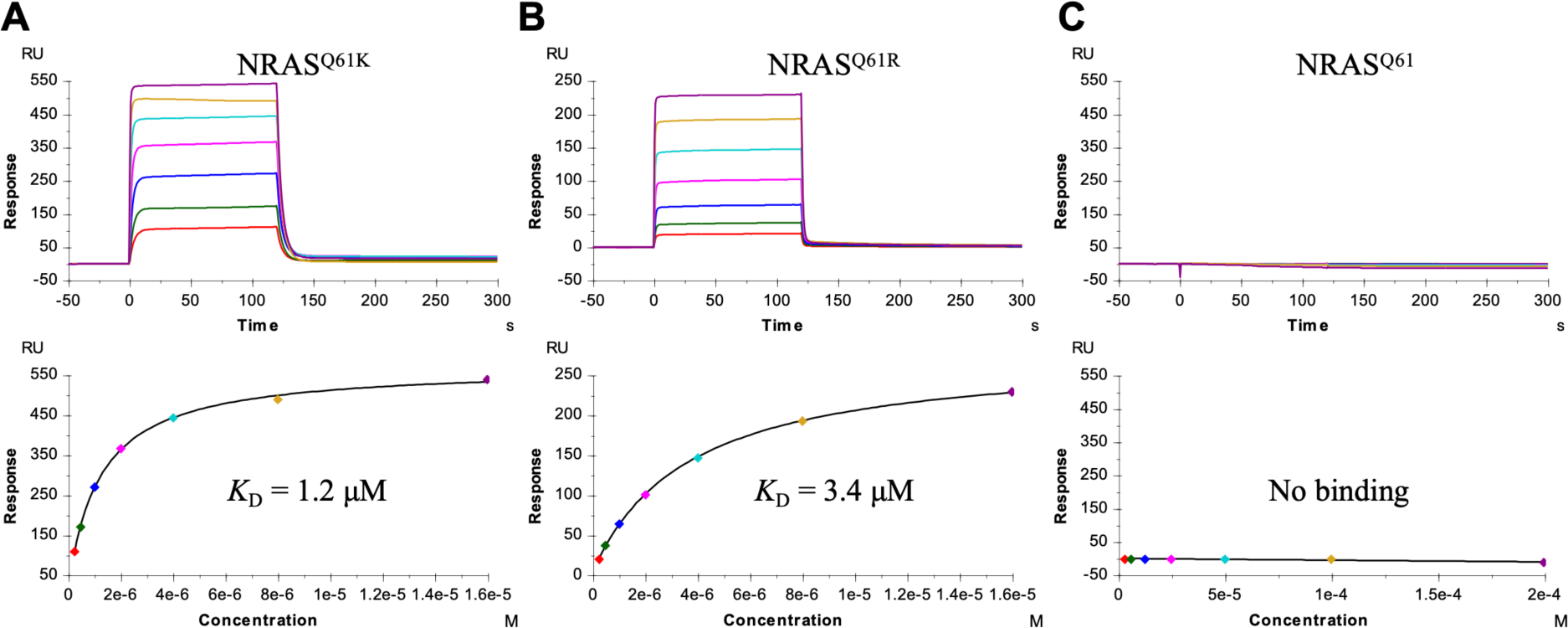
SPR analysis of TCR N17.1.2 binding to NRAS^Q61K^–HLA-A1, NRAS^Q61R^–HLA-A1, and NRAS–HLA-A1. (*A*) (upper) TCR N17.1.2 at concentrations of 0.25, 0.5, 1, 2, 4, 8, and 16 μM was injected over immobilized NRAS^Q61K^–HLA-A1 (1000 RU). (lower) Fitting curve for equilibrium binding that resulted in a *K*_D_ of 1.2 ± 0.1 μM. (*B*) (upper) TCR N17.1.2 at concentrations of 0.25, 0.5, 1, 2, 4, 8, and 16 μM was injected over immobilized NRAS^Q61R^–HLA-A1. (lower) Fitting curve for equilibrium binding that resulted in a *K*_D_ of 3.4 ± 0.2 μM. (*C*) (upper) TCR N17.1.2 at concentrations of 3.12, 6.25, 12.5, 25.5, 50, 100, and 200 μM was injected over immobilized NRAS^Q61^–HLA-A1. (lower) Fitting curve for equilibrium binding that showed no interaction.

### Overview of the N17.1.2–NRAS^Q61K^–HLA-A1 and N17.1.2–NRAS^Q61R^–HLA-A1 Complexes

To understand how TCR N17.1.2 discriminates between wild-type and mutant NRAS epitopes (**Fig. 1**), and how it recognizes both NRAS^Q61K^ and NRAS^Q61R^, we determined the structures of the N17.1.2–NRAS^Q61K^–HLA-A1 and N17.1.2–NRAS^Q61R^–HLA-A1 complexes to 2.10 Å and 2.26 Å resolution, respectively (***SI Appendix*, Table S1**) (**Fig. 2*A*, *D***). The interface between TCR and pMHC was in unambiguous electron density in both structures (***SI Appendix*, Fig. S1**). The root-mean-square difference (RMSD) in α-carbon positions for the two complexes is 0.3 Å, indicating very close similarity. TCR N17.1.2 docks over NRAS^Q61K^–HLA-A1 and NRAS^Q61R^– HLA-A1 in a canonical diagonal orientation, with Vα over the α2 helix of HLA-A1 and Vβ over the α1 helix, and a TCR–pMHC crossing angle (30) of 28° for both complexes (**Fig. 2*B*, *E***). The complexes are similar with respect to incident angle, which corresponds to the degree of tilt of TCR over pMHC (31): 5° for NRAS^Q61K^–HLA-A1 and 7° for NRAS^Q61R^–HLA-A1. The TCR N17.1.2 binding position is shifted toward the peptide N-terminus relative to most other TCR– pMHC structures; in comparison with a set of 82 reference TCR–pMHC class I complex structures, N17.1.2 has more peptide N-terminal shift than 79 (NRAS^Q61K^–HLA-A1 complex) and 80 (NRAS^Q61R^–HLA-A1 complex) structures (***SI Appendix*, Table S2**). As depicted by the footprint of TCR N17.1.2 on the pMHC surface (**Fig. 2*C*, *F***), N17.1.2 establishes contacts with the N-terminal half of the NRAS^Q61K^ or NRAS^Q61R^ peptide mainly through the CDR1α and CDR3α loops, whereas the CDR3β loop mostly contacts the C-terminal half.

**Fig. 2.**
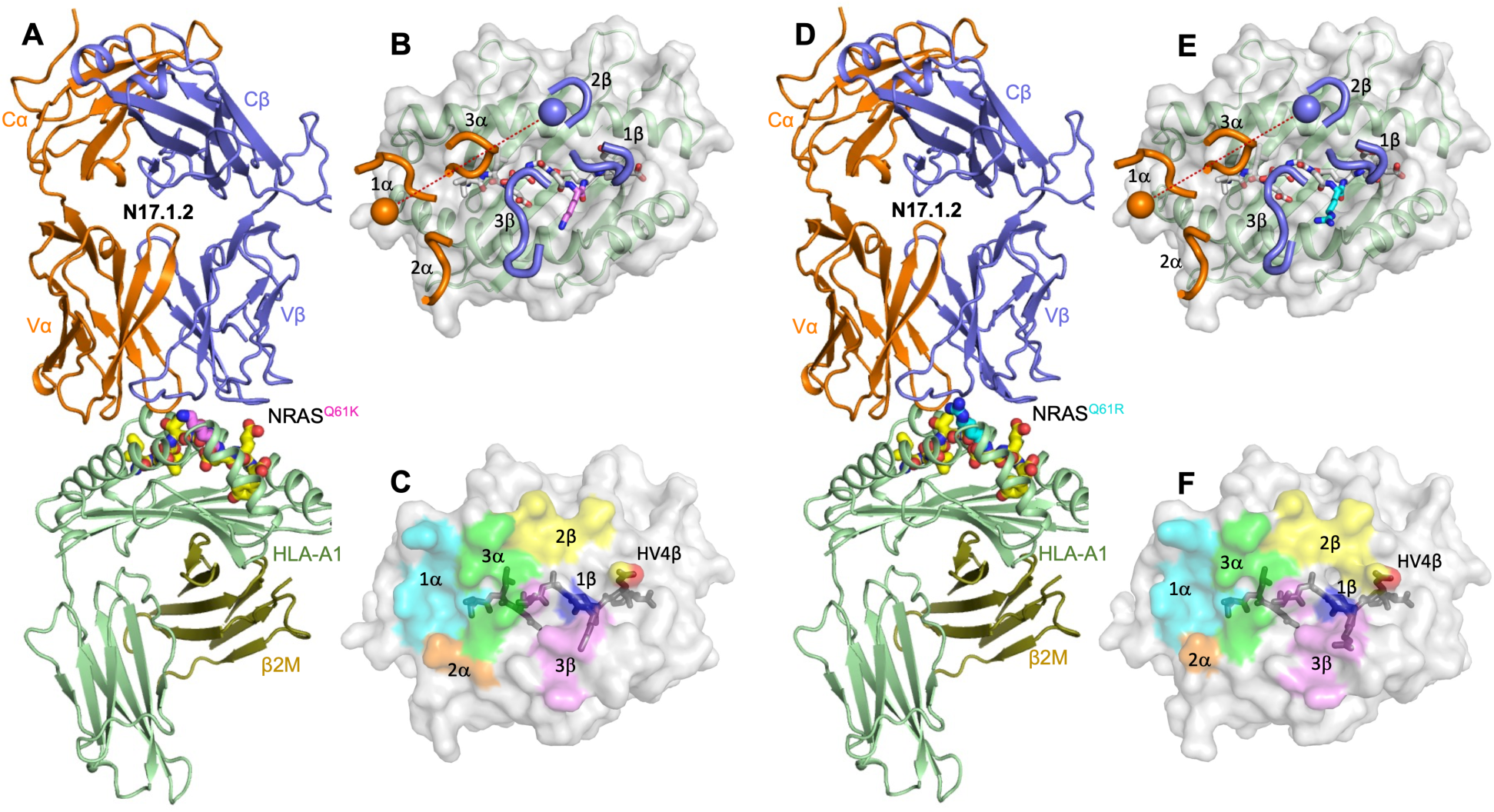
Structure of TCR N17.1.2 in complex with NRAS neoantigens. (*A*) Side view of the TCR N17.1.2–NRAS^Q61K^–HLA-A1 complex. (*B*) Positions of CDR loops of TCR N17.1.2 on NRAS^Q61K^–HLA-A1 (top view). CDRs of N17.1.2 are shown as numbered brown (CDR1α, CDR2α, and CDR3α) or blue (CDR1β, CDR2β, and CDR3β) loops. HLA-A1 is depicted as a gray surface and green cartoon. The NRAS^Q61K^ peptide is drawn in yellow in stick representation with the mutated P7 Lys residue in violet. The brown and blue spheres mark the positions of the conserved intrachain disulfide of the Vα and Vβ domains, respectively. The red dashed line indicates the crossing angle of TCR to pMHC. (*C*) Footprint of TCR N17.1.2 on NRAS^Q61K^–HLA-A1. The top of the MHC molecule is depicted as a gray surface. The areas contacted by individual CDR loops are color-coded: CDR1α, cyan; CDR2α, brown; CDR3α, green; CDR1β, blue; CDR2β, yellow; HV4β, red; CDR3β, violet. (*D*) Side view of the TCR N17.1.2–NRAS^Q61R^–HLA-A1 complex. (*E*) Positions of CDR loops of TCR N17.1.2 on NRAS^Q61R^–HLA-A1 (top view). The NRAS^Q61R^ peptide is drawn in yellow in stick representation with the mutated P7 Arg residue in cyan. (*F*) Footprint of TCR N17.1.2 on NRAS^Q61R^–HLA-A1.

### Vα Dominates Interactions with MHC

Of the total number of contacts (128) that TCR N17.1.2 makes with HLA-A1 in its complex with NRAS^Q61K^–HLA-A1, excluding the NRAS^Q61K^ peptide, CDR1α, CDR2α and CDR3α contribute 36%, 9%, and 26%, respectively, compared with 0%, 16% and 13% for CDR1β, CDR2β and CDR3β, respectively (**Table 1**) (**Fig. 3*E***). A very similar distribution of TCR contacts with MHC is observed in the N17.1.2–NRAS^Q61R^–HLA-A1 complex (**Fig. 3*F***). Hence, Vα dominates the interactions of N17.1.2 with MHC (90 of 128 contacts; 71%), with the germline-encoded CDR1α loop accounting for more of the binding interface (46 of 128 contacts; 36%) than any other CDR. TCR N17.1.2 makes substantially more interactions with the HLA-A1 α1 helix (**Fig. 3*A*, *B***) than the α2 helix (**Fig. 3*C*, *D***). These include a dense network of six hydrogen bonds linking Asp95α and Thr96α to Gln62H, Arg65H, and Asn66H of helix α1 (***SI Appendix*, Table S3**) (**Fig. 3*A*, *B***). In addition, Trp30α makes 22 van der Waals contacts and one hydrogen bond with Arg170H at the C-terminus of helix α2 to further anchor N17.1.2 to HLA-A1 (**Fig. 3*G***). Computational alanine scanning mutagenesis (32) showed eight HLA-A1 residues as possible hotspots for N17.1.2 binding in the NRAS^Q61K^–HLA-A1 complex, including residues Arg65H, Gln155H, Arg163H, and Arg170H; similar energies and hotspots were observed for the NRAS^Q61R^–HLA-A1 complex (***SI Appendix*, Table S4**).

**Fig. 3.**
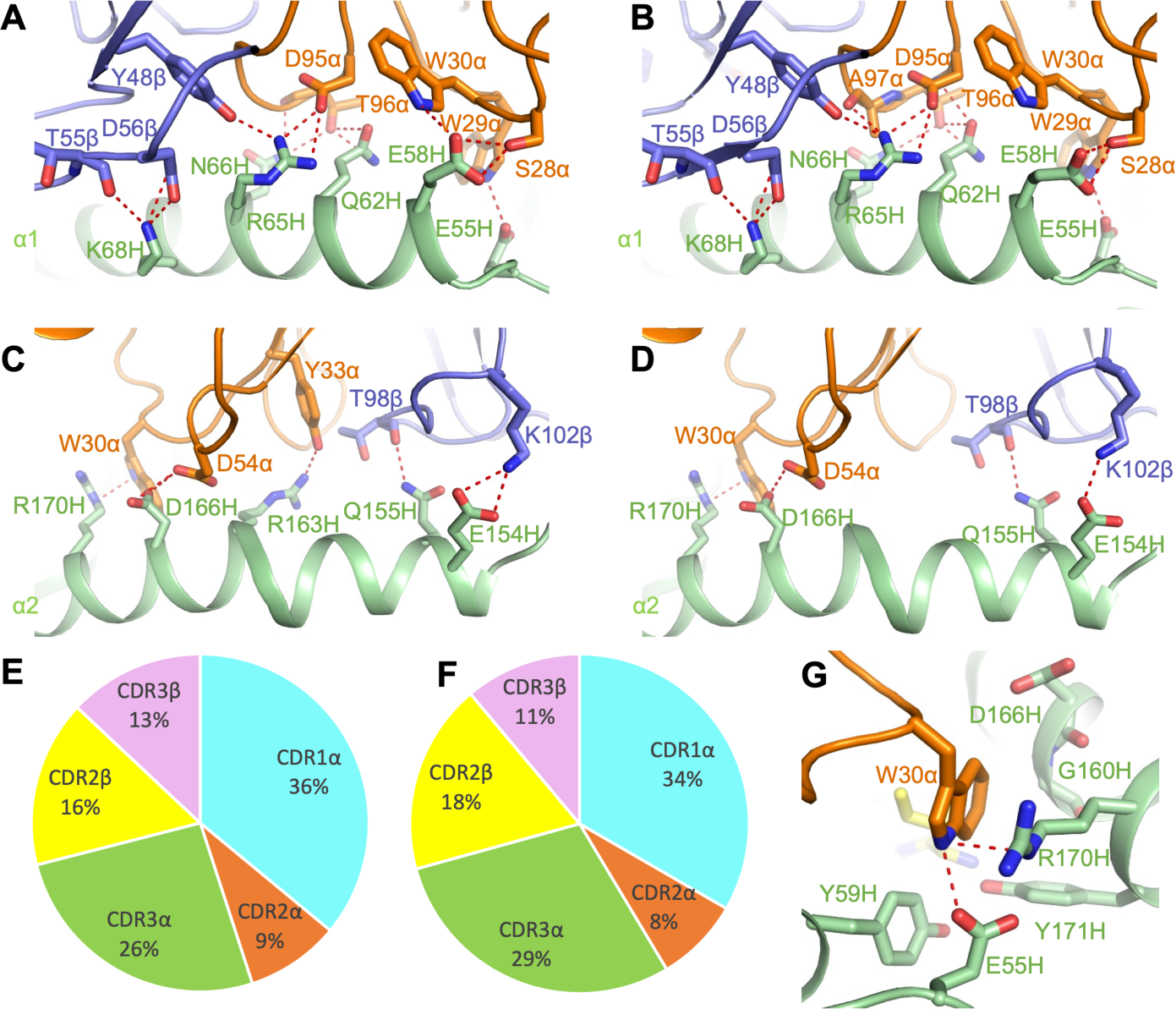
Interactions of TCR N17.1.2 with HLA-A1. (*A*) Interactions between N17.1.2 and the HLA-A1 α1 helix in the N17.1.2–NRAS^Q61K^–HLA-A1 complex. The side chains of contacting residues are drawn in stick representation with carbon atoms in brown (TCR α chain), blue (TCR β chain) or green (HLA-A1). Hydrogen bonds are indicated by red dashed lines. (*B*) Interactions between N17.1.2 and the HLA-A1 α1 helix in the N17.1.2–NRAS^Q61R^–HLA-A1 complex. (*C*) Interactions between N17.1.2 and the HLA-A1 α2 helix in the N17.1.2–NRAS^Q61K^–HLA-A1 complex. (*D*) Interactions between N17.1.2 and the HLA-A1 α2 helix in the N17.1.2–NRAS^Q61R^–HLA-A1 complex. (*E*) Pie chart showing percentage distribution of TCR N17.1.2 contacts to HLA-A1 according to CDR in the N17.1.2–NRAS^Q61K^–HLA-A1 complex. (f) Pie chart showing percentage distribution of TCR N17.1.2 contacts to HLA-A1 according to CDR in the N17.1.2–NRAS^Q61R^– HLA-A1 complex. (*F*) Close up view of interactions between Trp30α of N17.1.2 and Arg170H of HLA-A1.

**Table 1.**
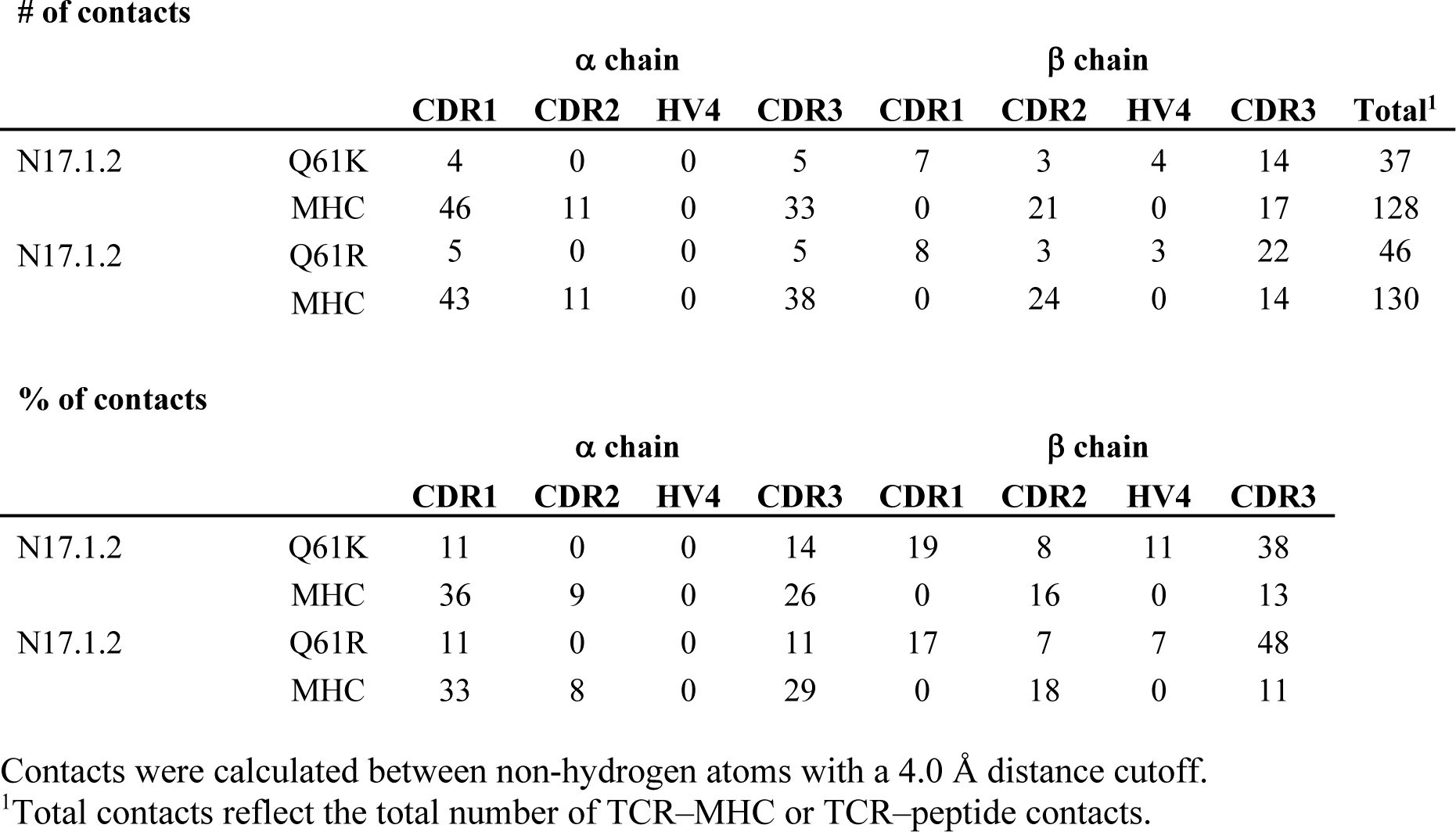
TCR CDR atomic contacts with NRAS^Q61K/R^ peptide and MHC.

| # of contacts | | $\alpha$ chain | | | | $\beta$ chain | | | | Total<sup>1</sup> |
| --- | --- | --- | --- | --- | --- | --- | --- | --- | --- | --- |
|  |  | CDR1 | CDR2 | HV4 | CDR3 | CDR1 | CDR2 | HV4 | CDR3 |  |
| N17.1.2 | Q61K | 4 | 0 | 0 | 5 | 7 | 3 | 4 | 14 | 37 |
|  | MHC | 46 | 11 | 0 | 33 | 0 | 21 | 0 | 17 | 128 |
| N17.1.2 | Q61R | 5 | 0 | 0 | 5 | 8 | 3 | 3 | 22 | 46 |
|  | MHC | 43 | 11 | 0 | 38 | 0 | 24 | 0 | 14 | 130 |

| % of contacts | | $\alpha$ chain | | | | $\beta$ chain | | | | |
| --- | --- | --- | --- | --- | --- | --- | --- | --- | --- | --- |
|  |  | CDR1 | CDR2 | HV4 | CDR3 | CDR1 | CDR2 | HV4 | CDR3 |  |
| N17.1.2 | Q61K | 11 | 0 | 0 | 14 | 19 | 8 | 11 | 38 |  |
|  | MHC | 36 | 9 | 0 | 26 | 0 | 16 | 0 | 13 |  |
| N17.1.2 | Q61R | 11 | 0 | 0 | 11 | 17 | 7 | 7 | 48 |  |
|  | MHC | 33 | 8 | 0 | 29 | 0 | 18 | 0 | 11 |  |

Contacts were calculated between non-hydrogen atoms with a 4.0 Å distance cutoff.
<sup>1</sup>Total contacts reflect the total number of TCR–MHC or TCR–peptide contacts.

### N17.1.2 Targets the NRAS^Q61K^ and NRAS^Q61R^ Mutations

In the solution structure of unbound mutant NRAS^Q61K^–HLA-A1 (20), the solvent-exposed side chains of P7 Lys and P9 Glu project away from the peptide-binding groove and present a prominent surface feature for potential interactions with TCR. Although the structure of wild-type NRAS–HLA-A1 is unknown, it likely differs from that of mutant NRAS^Q61K^–HLA-A1 only at P7 Lys, such that the structural differences which disclosed the naturally altered NRAS peptide to the T cells of cancer patients (15) are probably restricted to the mutation site at P7 and do not involve changes in peptide interactions with MHC. Similar considerations apply to the NRAS^Q61R^ neoepitope.

Upon binding NRAS^Q61K^–HLA-A1, TCR 17.1.2 buries 70% (273 Å^2^) of the peptide solvent-accessible surface, which is typical for TCR–pMHC complexes. However, TCR N17.1.2 makes relatively few contacts with peptide (37; 22%) compared to MHC (128; 78%) (**Table 1**). By contrast, three previously characterized TCRs (1a2, 12-6, and 38-10) specific for the p53^R175H^ neoepitope presented by HLA-A2 make many more contacts with peptide than does N17.1.2, both in absolute number (61, 64, and 82, respectively) and as a percentage of total interfacial contacts (50%, 41%, and 68%, respectively) (33). The N17.1.2 peptide contact percentage is also substantially lower than the median value among 82 reference TCR–pMHC class I structures (38%), although some complexes have comparable or lower peptide contact percentages (***SI Appendix*, Table S2**). Despite its relatively limited peptide contacts, TCR N17.1.2 is as specific for mutant versus wild-type NRAS (**Fig. 1**) as 1a2, 38-10, and 12-6 are for mutant versus wild-type p53 (33).

In both the N17.1.2–NRAS^Q61K^–HLA-A1 and N17.1.2–NRAS^Q61R^–HLA-A1 complexes, TCR 17.1.2 engages five residues of the mutant NRAS peptide (**Fig. 4*A*, *B***). However, the majority of interactions involve C-terminal residues P7 Lys/Arg and P9 Glu, whose protruding side chains are the most solvent-exposed: 13 of 29 van der Waals contacts and 7 of 8 hydrogen bonds for NRAS^Q61K^ and 21 of 40 van der Waals contacts and 6 of 7 hydrogen bonds for NRAS^Q61R^ (***SI Appendix*, Table S5**) (**Fig. 4*C*, *D***). These interactions primarily target the mutant P7 Lys/Arg residue, which forms a salt bridge with Glu103β of CDR3β in both complexes. The focus on P7 Lys/Arg suggests its functional importance for TCR binding. This conclusion is supported by binding energy calculations using Rosetta (32) to predict changes in TCR affinity upon alanine substitution of all peptide residues in the two complexes, in which P7 alanine substitutions led to substantial predicted loss of binding in both interfaces (***SI Appendix*,Table S6**). Other peptide residues identified as hotspots by Rosetta were P1 Ile (for NRAS^Q61K^, close to hotspot cutoff for NRAS^Q61R^), P4 Thr, and P9 Glu, indicating productive engagement of accessible side chains throughout the peptide by the TCR N17.1.2.

**Fig. 4.**
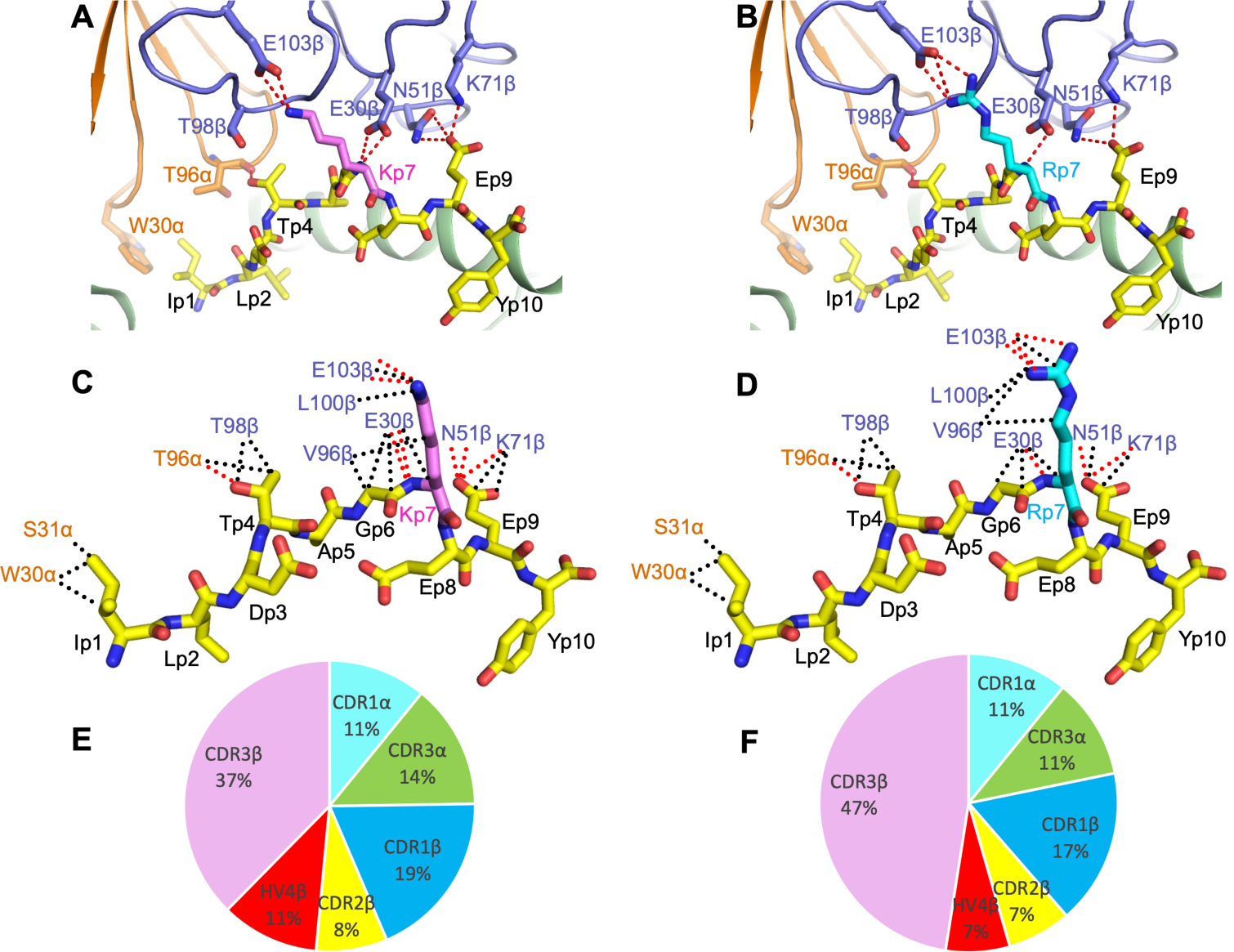
Interactions of TCR N17.1.2 with NRAS neoantigens. (*A*) Interactions between N17.1.2 and the NRAS^Q61K^ peptide. The side chains of contacting residues are shown in stick representation with carbon atoms in brown (TCR α chain), blue (TCR β chain), yellow (NRAS^Q61K^), or violet (mutated P7 Lys). Peptide residues are identified by one-letter amino acid designation followed by position (p) number. Hydrogen bonds are indicated by red dashed lines. (*B*) Interactions between N17.1.2 and the NRAS^Q61R^ peptide. The mutated P7 Arg residue is shown in cyan. (*C*) Schematic representation of N17.1.2–NRAS^Q61K^ interactions. Hydrogen bonds are red dotted lines and van der Waals contacts are black dotted lines. For clarity, not all van der Waals contacts are shown. (*D*) Schematic representation of N17.1.2–NRAS^Q61R^ interactions. (*E*) Pie chart showing percentage distribution of TCR N17.1.2 contacts to NRAS^Q61K^ according to CDR. (*F*) Pie chart showing percentage distribution of TCR N17.1.2 contacts to NRAS^Q61R^ according to CDR.

TCR N17.1.2 discriminates between mutant and wild-type NRAS by minimizing interactions with the N-terminal and central portions of NRAS^Q61K/R^, which are identical in mutant and wild-type peptides, and instead focusing on the P7 Lys/Arg mutation in the C-terminal portion. Of the 37 total contacts that N17.1.2 establishes with NRAS^Q61K^, the bulk (28; 76%) are mediated by Vβ (**Table 1**). Similarly, Vβ accounts for 36 of 46 total contacts (79%) with NRAS^Q61R^. Thus, whereas Vα dominates MHC recognition, Vβ dominates peptide recognition in both N17.1.2– NRAS^Q61K^–HLA-A1 and N17.1.2–NRAS^Q61R^–HLA-A1 complexes, with CDR3β making a considerably greater contribution than any other CDR (37% of contacts to NRAS^Q61K^; 47% of contacts to NRAS^Q61R^) (**Table 1**) (**Fig. 4*E*, *F***). In particular, Glu103β of the long CDR3β loop forms a salt bridge with P7 Lys/Arg (**Fig. 4*A*, *B***). Wild-type P7 Gln would be unable to replicate this key interaction, thereby explaining, at least in part, the exquisite specificity of TCR N17.1.2 for NRAS^Q61K/R^ (**Fig. 1**).

To computationally assess the energetic effect of replacing P7 Lys/Arg by Gln, which corresponds to reversion to the wild-type NRAS peptide, we performed in silico mutagenesis using Rosetta (32). Both reversion substitutions were predicted to be substantially disruptive for TCR binding, with ΔΔ*G* values of 1.2 and 1.7 Rosetta Energy Units, comparable to kcal/mol, for NRAS^K61Q^ and NRAS^R61Q^, respectively (***SI Appendix*, Table S6**).

### Limited Conformational Changes in TCR and pMHC upon Binding

To assess possible conformational selection or induced fit underlying N17.1.2 recognition of the NRAS^Q61K^–HLA-A1 pMHC ligand, we compared the structure of the NRAS^Q61K^ peptide in the N17.1.2-bound complex with the corresponding peptide in the previously determined NMR structure of unbound NRAS^Q61K^–HLA-A1 (PDB code 6MPP) (20). RMSDs calculated at each peptide position between TCR-bound and unbound peptide conformations, for each of 10 unbound NMR structure models, indicated relatively limited binding conformational changes for backbone atoms (***SI Appendix*, Fig. S2*A***), generally 1 Å binding RMSD on average across residues and unbound NMR models. More variability was observed when taking side chain atoms into account (all-atom RMSDs; ***SI Appendix*, Fig. S2B**), including at position P7 (mutated NRAS residue Lys61), however certain unbound conformations, including NMR model 7, showed relatively low all-atom unbound-bound RMSD at that position. Overall, unbound pMHC NMR model 7 has a backbone RMSD of 1.1 Å and all-atom RMSD of 1.4 Å in comparison with the N17.1.2-bound peptide. Given this approximate representation of the N17.1.2-bound state among the unbound NMR models, TCR N17.1.2 recognition of NRAS^Q61K^–HLA-A1 appears to be largely a conformational selection scenario, versus induced fit.

To identify possible ligand-induced conformational changes in TCR N17.1.2, we determined its structure in unbound form to 3.50 Å resolution (***SI Appendix*, Table S1**). Superposition of the VαVβ domains of free N17.1.2 onto those of N17.1.2 in complex with NRAS^Q61K^–HLA-A1 or NRAS^Q61R^–HLA-A1 revealed no notable structural differences in any of the six CDR loops, including CDR3α and CDR3β, which frequently undergo significant shifts upon engaging pMHC (***SI Appendix*, Fig. S3**). Structural differences are limited to small side chain movements. Overall, the CDR loops of unbound N17.1.2 have a backbone RMSD of 0.53 Å and an all-atom RMSD of 0.62 Å compared with the TCR bound to NRAS^Q61K^–HLA-A1 (0.50 Å and 0.61 Å, respectively, compared with the TCR bound to NRAS^Q61R^–HLA-A1). Hence, N17.1.2 behaves essentially as a rigid body in binding pMHC.

### AlphaFold Structure Predictions and Comparison with Crystal Structures

To test the capability of AlphaFold (22) to model previously unseen TCR–pMHC complexes, we used AlphaFold v.2.3 in the TCRmodel2 framework (25) to generate models of the N17.1.2– NRAS^Q61K^–HLA-A1 and N17.1.2–NRAS^Q61R^–HLA-A1 complexes from sequence, in comparison with AlphaFold2.3 (**Table 2**). Additionally, we tested the recently released AlphaFold3 algorithm, which includes a generalized representation of proteins and other molecules and was reported to improve performance of antibody–antigen modeling versus AlphaFold2.3 (34). Models were ranked based on AlphaFold’s standard model confidence score, and interface pLDDT (I-pLDDT) score was calculated as an additional confidence metric for each model, based on AlphaFold’s residue-level confidence scores (pLDDT) for residues at the TCR–pMHC interface. As default TCRmodel2 showed low-to-moderate model confidence scores and low model accuracies for top-ranked models for each complex, we generated additional models of each complex (1000 TCRmodel2 models for each complex, versus 5 models for default TCRmodel2), and assessed the top-ranked model scores and accuracies from that expanded set (**Table 2**). This approach is similar to a previously described AlphaFold massive sampling approach (35), which we recently found was effective for improving antibody–antigen modeling success (36), and it is also reflective of the strategy used by the AlphaFold3 team for antibody-antigen modeling (34). Such approaches are based on the principle that AlphaFold’s scoring can be generally accurate, but for challenging complexes, additional models need to be generated stochastically to enable sampling of near-native conformations. Top-ranked models from that protocol had higher AlphaFold confidence scores and were found to be accurate with respect to the X-ray complex structures (high accuracy, based on CAPRI criteria (37)), while AlphaFold3 models notably had similar, although slightly lower, accuracy (**Table 2**). Comparison of the TCRmodel2-modeled structures with experimental density maps for N17.1.2–NRAS^Q61K^–HLA-A1 (**Fig. 5**) and N17.1.2–NRAS^Q61R^–HLA-A1 (***SI Appendix*, Fig. S4**) shows that the models largely reflect key interface residue positions and interactions, however the experimentally determined structures fit the density maps better for some residues. We also assessed TCR–peptide polar and charged contacts for the models, and found that the model-derived interface polar contacts were in close agreement with those based on the X-ray structures (***SI Appendix*, Table S7**). These include four of eight TCR–peptide hydrogen bonds in the N17.1.2–NRAS^Q61K^–HLA-A1 complex and four of seven TCR–peptide hydrogen bonds in the N17.1.2–NRAS^Q61R^–HLA-A1 complex.

**Fig. 5.**
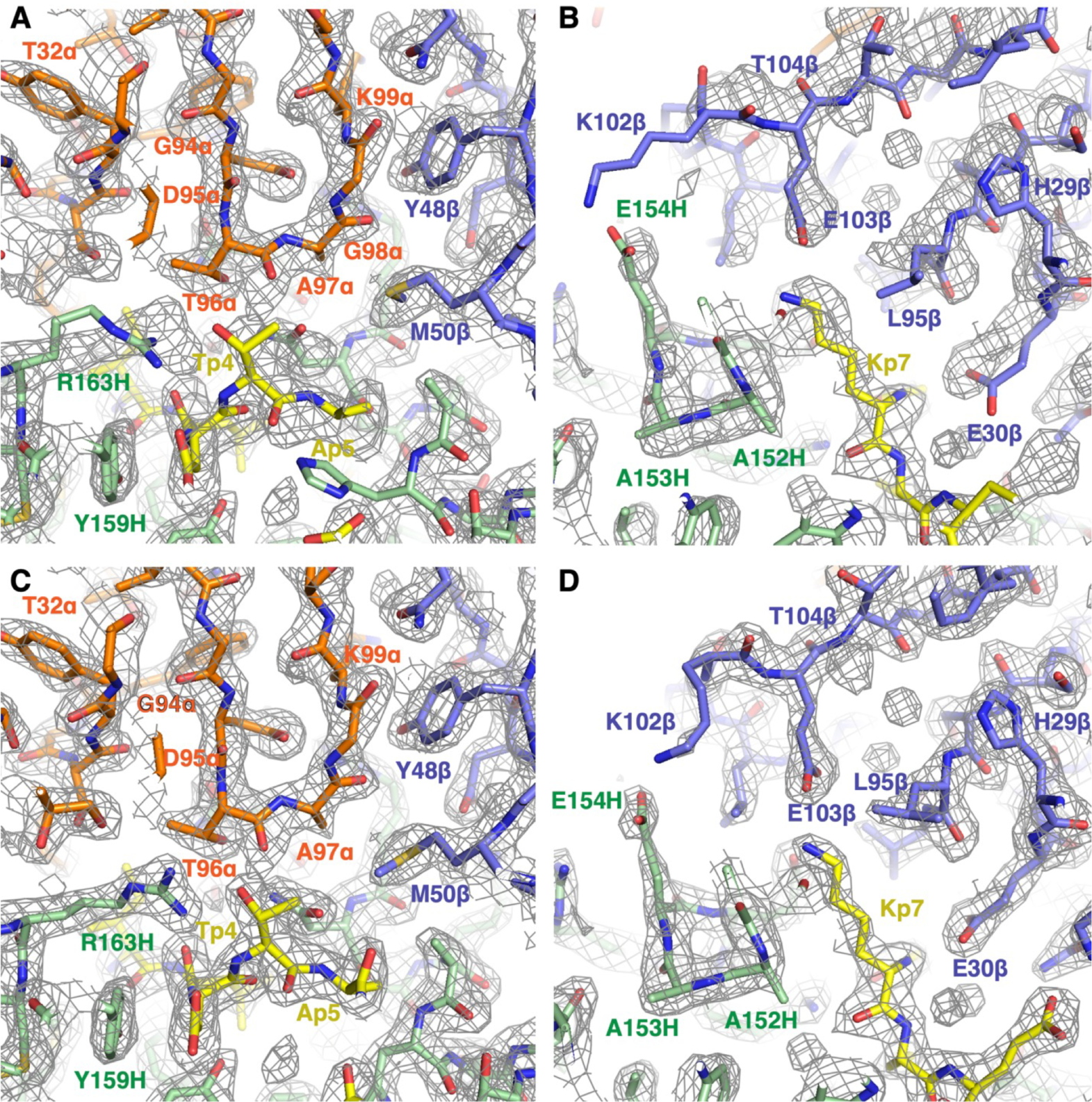
Comparison of AlphaFold prediction of TCR N17.1.2–NRAS^Q61K^–HLA-A1 complex generated by TCRmodel2 with crystallographic density map and structures. (*A*) AlphaFold prediction of N17.1.2–NRAS^Q61K^–HLA-A1 complex compared with experimental electron density map in the region of CDR3α (Vα, orange; Vβ, blue; NRAS^Q61K^ peptide, yellow; HLA-A1, green). (*B*) AlphaFold prediction of N17.1.2–NRAS^Q61K^–HLA-A1 complex compared with electron density map in the region of CDR3β. (*C*) Model of N17.1.2–NRAS^Q61K^–HLA-A1 complex built into electron density map in the region of CDR3α (Vα, orange; Vβ, blue; NRAS^Q61K^ peptide, yellow; HLA-A1, green). (*D*) Model of N17.1.2–NRAS^Q61K^–HLA-A1 complex built into electron density map in the region of CDR3β.

**Table 2.** N17.1.2–NRAS^Q61K/R^–HLA-A1 AlphaFold and TCRmodel2 model scores and accuracies.

| Complex | Protocol<br>(# models)<sup>1</sup> | Model<br>Confidence<sup>2</sup> | I-pLDDT<sup>2</sup> | I-RMSD<br>(Å)<sup>3</sup> | L-RMSD<br>(Å)<sup>3</sup> | CAPRI<br>Accuracy<sup>3</sup> | DockQ<br>Score<sup>3</sup> |
| --- | --- | --- | --- | --- | --- | --- | --- |
| N17.1.2-NRAS<sup>Q61K</sup>-<br>HLA-A1 | AlphaFold2<br>(25) | 0.88 | 81.62 | 4.70 | 10.20 | Incorrect | 0.250 |
| N17.1.2-NRAS<sup>Q61R</sup>-<br>HLA-A1 | AlphaFold2<br>(25) | 0.89 | 82.21 | 4.91 | 10.73 | Incorrect | 0.244 |
| N17.1.2-NRAS<sup>Q61K</sup>-<br>HLA-A1 | TCRmodel2<br>(5) | 0.87 | 79.29 | 4.11 | 8.08 | Acceptable | 0.284 |
| N17.1.2-NRAS<sup>Q61R</sup>-<br>HLA-A1 | TCRmodel2<br>(5) | 0.88 | 84.25 | 7.10 | 16.63 | Incorrect | 0.135 |
| **N17.1.2-NRAS<sup>Q61K</sup>-**<br>**HLA-A1** | **TCRmodel2**<br>**(1000)** | **0.92** | **93.5** | **0.88** | **2.92** | **High** | **0.823** |
| **N17.1.2-NRAS<sup>Q61R</sup>-**<br>**HLA-A1** | **TCRmodel2**<br>**(1000)** | **0.92** | **94.02** | **0.85** | **2.39** | **High** | **0.829** |
| N17.1.2-NRAS<sup>Q61K</sup>-<br>HLA-A1 | AlphaFold3 | 0.94 | 89.05 | 1.31 | 3.99 | Medium | 0.731 |
| N17.1.2-NRAS<sup>Q61R</sup>-<br>HLA-A1 | AlphaFold3 | 0.94 | 88.30 | 1.14 | 2.26 | Medium | 0.774 |

<sup>1</sup>AlphaFold-based modeling protocol, either default AlphaFold2.3 (AlphaFold2) (22), TCRmodel2 (25), or AlphaFold3 (34). AlphaFold2.3 generated 25 models per complex, based on its default setting, and TCRmodel2 was used to generate 5 (default) or 1000 models per complex, as noted. AlphaFold3 was run on its web server which generated 5 models per complex. The top-ranked model from each set was selected based on AlphaFold model confidence score and assessed for accuracy.
<sup>2</sup>AlphaFold model confidence and interface pLDDT (I-pLDDT) confidence scores for top-ranked model from the given protocol. For AlphaFold3, the provided “ranking\_score”, which includes a slight modification of the AlphaFold2.3 model confidence score calculation, is shown for model confidence.
<sup>3</sup>Model accuracy metrics based on comparison with the X-ray structure of the corresponding complex and computed by the DockQ program (51). Shown are interface RMSD (I-RMSD), ligand RMSD (L-RMSD), CAPRI accuracy, and DockQ score. Models with High CAPRI accuracy shown in bold.

After noting the capability of AlphaFold and TCRmodel2 to generate accurate models of the N17.1.2 complexes, we used it to generate models of complexes for four additional TCRs that were experimentally confirmed to target NRAS^Q61K^–HLA-A1 (15). Given that high AlphaFold model confidence scores are associated with accurate models, both for the N17.1.2 complexes described here and based on previous benchmarking (25), we assessed model accuracies using a benchmark set of 20 TCR–pMHC complexes from our previous study and the TCRmodel2 protocol used here (1000 models per complex), to provide additional information on expected accuracies above defined confidence score thresholds that can inform predictive modeling. We found that individual model confidence and I-pLDDT cutoffs of 0.875 and 87.5 led to moderate rates of high accuracy models (with I-pLDDT performing better), while the combination of those two cutoffs enriched the proportion of high accuracy models further, with 75% high accuracy and the remaining models with medium accuracy (***SI Appendix*, Fig. S5**). Thus, we used the combination of score cutoffs (model confidence > 0.875, I-pLDDT > 0.875) to identify likely accurate models in our predictive modeling. The model confidence cutoff is more strict than noted previously for TCRmodel2 (model confidence ≥ 0.84) (25); this difference is due to the emphasis here on achieving models closer to native (high CAPRI accuracy), versus slightly more permissive medium or high CAPRI accuracy in the previous study. Surprisingly, we found that the predictive models for the four NRAS^Q61K^-specific TCRs did not yield sufficiently high model confidence scores, using TCRmodel2 (1000 models), AlphaFold2.3 (200 models), and AlphaFold3 (5 models) (***SI Appendix*, Table S8**), despite the capability of the TCRmodel2 and AlphaFold3 to achieve relatively high confidence scores for the N17.1.2 complexes. Of note, the other NRAS^Q61K^-specific TCRs have distinct sequences, and therefore likely distinct modes of pMHC engagement, from N17.1.2. It is possible that more extensive sampling strategies in TCRmodel2 or AlphaFold3 would lead to higher confidences scores and predictive success for those complexes. Additionally, it is possible that accurate models were predicted, but with unexpectedly low scores (false negatives). Regardless, this indicates that improved sampling and/or scoring is needed to systematically model unknown TCR-pMHC complex structures.

## Discussion

Structural studies of mutated self-antigens have provided insights into how different a neoantigen must be from its wild-type parent for it to induce a T cell response and into the multiple mechanisms TCRs employ to detect cancer neoantigens (21). In some cases, the mutation improves antigen presentation by strengthening peptide–MHC binding. For example, a glycine-to-aspartate mutation in the KRAS^G12D^ neoepitope enables P3 Asp (the mutant residue) to make a stabilizing salt bridge with Arg156 on the α2 helix of HLA-C that cannot form with P3 Gly (the wild-type residue) (38). In other cases, the mutation does not affect peptide–MHC binding or antigen presentation, but instead increases affinity for TCR, either through direct contacts with TCR or through indirect strategies not requiring direct TCR contacts with the mutation (21). For example, an arginine-to-histidine mutation in the p53^R175H^ neoepitope enables P8 His to make multiple hydrogen bonds with TCRs specific for p53^R175H^–HLA-A2 that cannot form with P8 Arg (33). Similarly, TCR N17.1.2 detects the NRAS^Q61K^ and NRAS^Q61R^ neoepitopes by directly targeting the P7 Lys/Arg mutation through formation of a critical salt bridge with Glu103β of CDR3β that cannot be made by P7 Gln.

In this study, we found that AlphaFold was able to generate accurate models of TCR N17.1.2 in complex with its peptide–MHC targets, however this success was only after increased sampling in TCRmodel2 which led to high model confidence scores. This underscores the utility of model confidence scores, which were shown previously for TCR–pMHC modeling (25) and more recently for antibody–antigen modeling (36) to be indicators of accurate models. Notably, the confidence score thresholds identified and used here are higher here than in Yin et al. (25), due to the emphasis here on high accuracy (based on CAPRI criteria (37)) models that can reflect more details from the experimentally determined interfaces. In contrast with this study, others have recently observed that AlphaFold failed to accurately model four complexes of antibodies with peptide–MHC class I targets (39). The lack of success observed in that study could stem from various factors including limited sampling in AlphaFold and antigen size; protein complex size previously showed some association with lower modeling success in AlphaFold (27, 36). Additionally, TCR–pMHC complexes are generally more successfully modeled in AlphaFold than antibody–antigen complexes as shown through benchmarking (25, 36), possibly due to the potential co-evolutionary signal between TCR and MHC protein sequences in AlphaFold’s database and multiple sequence alignments, or general TCR–pMHC docking topology information learned by AlphaFold during its training, which could reduce its complex modeling to a more local structural search. Notably, the AlphaFold3 model accuracies were higher than for AlphaFold2.3 for N17.1.2 complex modeling and nearly as accurate as TCRmodel2 with additional sampling, indicating that AlphaFold3 may generally improve TCR–pMHC complex modeling accuracy over AlphaFold2.3, as the authors noted for antibody–antigen complex modeling (34). As observed in this study with TCRmodel2, additional sampling was found to improve AlphaFold3 accuracy for antibody-antigen modeling (34); it is currently not practical to run such a protocol (1000 seeds, corresponding to 5000 models generated (34)) through the current AlphaFold3 web server interface, but it is possible that such an approach would improve AlphaFold3 accuracy for N17.1.2 or other TCR-pMHC complexes.

The reason underlying the successful modeling of the N17.1.2 complexes, versus the other NRAS^Q61K^ TCR complexes considered in this study, is not yet clear. We observed relatively limited conformational changes in TCR N17.1.2 and its CDR loops associated with binding, which would result in it being an easier target for traditional modeling approaches (e.g. TCRFlexDock (40)), however previous benchmarking has shown that AlphaFold’s complex modeling performance is not impacted by the degree of binding conformational changes (27). However, this has not been assessed directly for TCRs or antibodies, due in part to limited sets of recently determined unbound antibody and TCR structures. Additionally, it is possible that at least some of the other NRAS^Q61K^ TCR complexes were modeled accurately, but those models not appropriately identified by the model confidence scores and thresholds used in this study (false negative). Fine-tuning of the AlphaFold model, or additional sampling or scoring strategies in AlphaFold2, AlphaFold3, and other deep learning structure prediction methods, may enable improved predictive success for TCR–pMHC complexes. Such advances would be applicable toward the major challenge of modeling and mapping of T cell specificities on a large scale (26).

## Materials and Methods

### Protein Preparation

The isolation and characterization of NRAS^Q61K^-specific TCR N17.1.2 from melanoma patients was described previously (15). Soluble TCR N17.1.2 for affinity measurements and structure determination was produced by *in vitro* folding from inclusion bodies expressed in *Escherichia coli*. Codon-optimized genes encoding the TCR α and β chains (residues 1–206) and 1–246, respectively) were synthesized and cloned into the expression vector pET22b (GenScript). An interchain disulfide (CαCys160–CβCys173) was engineered to increase the folding yield of TCR αβ heterodimers (41). The mutated α and β chains were expressed separately as inclusion bodies in BL21(DE3) *E. coli* cells (Agilent Technologies). Bacteria were grown at 37 °C in LB medium to OD_600_ = 0.6–0.8 and induced with 1 mM isopropyl-β-D-thiogalactoside. After incubation for 3 h, the bacteria were harvested by centrifugation and resuspended in 50 mM Tris-HCl (pH 8.0) containing 0.1 M NaCl and 2 mM EDTA. Cells were disrupted by sonication. Inclusion bodies were washed with 50 mM Tris-HCl (pH 8.0) and 5% (v/v) Triton X-100, then dissolved in 8 M urea, 50 mM Tris-HCl (pH 8.0), 10 mM EDTA, and 10 mM DTT. For *in vitro* folding, the TCR α (45 mg) and β (35 mg) chains were mixed and diluted into 1 liter folding buffer containing 5 M urea, 0.4 M L-arginine-HCl, 100 mM Tris-HCl (pH 8.0), 3.7 mM cystamine, and 6.6 mM cysteamine. After dialysis against 10 mM Tris-HCl (pH 8.0) for 72 h at 4 °C (buffer swapped at 48 h), the folding mixture was concentrated 20-fold and dialyzed against 50 mM MES buffer (pH 6.0) to precipitate misfolded protein. The supernatant was dialyzed overnight at 4 °C against 20 mM Tris-HCl (pH 8.0), 20 mM NaCl. Disulfide-linked TCR N17.1.2 was purified using sequential Superdex 200 (20 mM Tris-HCl (pH 8.0), 20 mM NaCl) and Mono Q (20 mM Tris-HCl (pH 8.0), 0–1.0 M NaCl gradient) FPLC columns (GE Healthcare).

Soluble HLA-A1 loaded with NRAS^Q61K^ peptide (ILDTAG**K**EEY), NRAS^Q61R^ peptide (ILDTAG**R**EEY) peptide, or wild-type NRAS peptide (ILDTAG**Q**EEY) peptide was prepared by *in vitro* folding of *E. coli* inclusion bodies as described (33). Correctly folded NRAS^Q61K^–HLA-A1, NRAS^Q61R^–HLA-A1, and NRAS–HLA-A1 complexes were purified using consecutive Superdex 200 (20 mM Tris-HCl (pH 8.0), 20 mM NaCl) and Mono Q columns (20 mM Tris-HCl (pH 8.0), 0–1.0 M NaCl gradient). To produce biotinylated HLA-A1, a C-terminal tag (GGGLNDIFEAQKIEWHE) was attached to the HLA-A*01:01 heavy chain. Biotinylation was carried out with BirA biotin ligase (Avidity). Biotinylated proteins were separated from excess biotin with a Superdex 200 column (20 mM Tris-HCl (pH 8.0), 20 mM NaCl).

### Crystallization and Data Collection

For crystallization of TCR–pMHC complexes, TCR N17.1.2 was mixed with NRAS^Q61K^–HLA-A1 or NRAS^Q61R^–HLA-A1 in a 1:1 ratio at a concentration of 10 mg/ml. Crystals were obtained at room temperature by vapor diffusion in hanging drops. The N17.1.2–NRAS^Q61K^–HLA-A1 complex crystallized in 8% (w/v) Tacsimate (pH 7.0) and 20% (w/v) polyethylene glycol (PEG) 4000 (Hampton Research). Crystals of the N17.1.2–NRAS^Q61R^–HLA-A1 complex grew in 0.2 M ammonium citrate tribasic (pH 7.4) and 16% (w/v) PEG 3350 by seeding. Unbound TCR N17.1.2 (8 mg/ml) crystallized in 0.2 M magnesium chloride, 0.2 M sodium nitrate, and 14–18% (w/v) PEG 3350. Before data collection, all crystals were cryoprotected with 20% (w/v) glycerol and flash-cooled. X-ray diffraction data were collected at beamline 23-ID-B of the Advanced Photon Source, Argonne National Laboratory. Diffraction data were indexed, integrated, and scaled using the program HKL2000 (42). Data collection statistics are shown in ***SI Appendix*, Table S1**.

### Structure Determination and Refinement

Before structure determination and refinement, all data reductions were performed using the CCP4 software suite (43). Structures were determined by molecular replacement with the program Phaser (44) and refined with Phenix (45). The models were further refined by manual model building with Coot (46) based on 2*F*_o_ – *F*_c_ and *F*_o_ – *F*_c_ maps. The γ chain of γδ TCR AB18.1 (PDB accession code 4NDM) (47), the β chain of p53^R175H^-specific TCR 1a2 (6VQO) (33), and NRAS^Q61K^–HLA-A1 (6MPP) (20) with the CDRs and peptide removed were used as search models to determine the orientation and position of the N17.1.2– NRAS^Q61K^–HLA-A1 complex. The orientation and position parameters of the N17.1.2– NRAS^Q61R^–HLA-A1 complex were obtained using the coordinates of the N17.1.2–NRAS^Q61K^– HLA-A1 complex as a search model. The TCR component of the N17.1.2–NRAS^Q61K^–HLA-A1 complex was used a search model to determine the coordinates of unbound N17.1.2. Refinement statistics are summarized in ***SI Appendix*, Table S1**. Contact residues were identified with the CONTACT program (43) and were defined as residues containing an atom 4.0 Å or less from a residue of the binding partner. The PyMOL program (https://pymol.org/) was used to prepare figures.

### Surface Plasmon Resonance Analysis

The interaction of TCR N17.1.2 with NRAS^Q61K^–HLA-A1, NRAS^Q61R^–HLA-A1, and wild-type NRAS–HLA-A1 was assessed by surface plasmon resonance (SPR) using a BIAcore T100 biosensor at 25 °C. Biotinylated NRAS^Q61K^–HLA-A1, NRAS^Q61R^–HLA-A1, or NRAS–HLA-A1 ligand was immobilized on a streptavidin-coated BIAcore SA chip (GE Healthcare) at approximately 1000 resonance units (RU). The remaining streptavidin sites were blocked with 20 μM biotin solution. An additional flow cell was injected with free biotin alone to serve as a blank control. For analysis of TCR binding, solutions containing different concentrations of N17.1.2 were flowed sequentially (50 μl/min, 600 s for dissociation) over chips immobilized with NRAS^Q61K^–HLA-A1, NRAS^Q61R^–HLA-A1, or NRAS–HLA-A1 ligand, or the blank. Dissociation constants (*K*_D_s) were calculated by fitting equilibrium and kinetic data to a 1:1 binding model using BIAevaluation 3.1 software.

### Computational Sequence and Structural Analysis

Calculation of TCR–pMHC incident and crossing angles was performed using a previously developed program that is available on the TCR3d database (31), and calculation of TCR position over peptide groove axis was performed using a Perl script as previously described (33). Rosetta v.2.3 was used to perform computational alanine scanning and model other point substitutions to calculate binding affinity changes (ΔΔ*G*s) using the interface mode (“-interface” flag) (32, 48), using default parameters except for extra side chain rotamers (“-extrachi_cutoff 1 -ex1 -ex2 -ex3” flags). Structures were pre-processed using the FastRelax protocol (49, 50) in Rosetta 3 (weekly release 2021.38) prior to computational mutagenesis, with coordinate constraints enabled (“-relax:constrain_relax_to_start_coords” flag).

### Structure Prediction

TCR–pMHC complexes were modeled from sequence using AlphaFold2 (22), AlphaFold3 (34), and TCRmodel2 (25), with TCR sequences trimmed to the Vα and Vβ domains, and HLA-A1 sequence was trimmed to α1 and α2 domains, for modeling efficiency. AlphaFold (v2.3.0) was obtained from Github (https://github.com/google-deepmind/alphafold) in February 2023 and installed on a local cluster. Either 25 or 200 predictions per complex were generated, with structural refinement performed on the top-ranked prediction, ranked by model confidence score. TCRmodel2 predictions were generated on a local cluster. To generate 1,000 predictions per complex, the “num_predictions_per_model” parameter was set to 200. Predictions were ranked by model confidence score, and structure refinement was performed on the top-ranked prediction. AlphaFold3 (34) was run through its public web server (https://www.alphafoldserver.com), which generated five models per complex, ranked by AlphaFold3 ranking score.

Interface-pLDDT (I-pLDDT) score was calculated as average pLDDT score of all interface residues, defined as residues with any atoms that are within 4 Å of the binding partner. Accuracy metrics of modeled structures with respect to X-ray structures were calculated by the DockQ program (51).

## Data Availability

Atomic coordinates and structure factors have been deposited in the Protein Data Bank under accession codes 8YIV (N17.1.2–NRAS^Q61K^–HLA-A1), 8YJ2 (N17.1.2– NRAS^Q61R^–HLA-A1), and 8YJ3 (N17.1.2).

## ACKNOWLEDGMENTS

This work was supported by National Institutes of Health Grants GM144083 (to B.G.P.) and AI129893 (to R.A.M.), National Natural Science Foundation of China Grants 32270995 (to D.W.) and 32100985 (to D.W.), Outstanding Youth Fund of Hunan Provincial Natural Science Foundation 2023JJ10034 (to D.W.), the Science and Technology Innovation Program of Hunan Province Grant 2022RC1209 (to D.W.), and by the National Cancer Institute-University of Maryland Partnership (to R.Y., B.G.P., and P.F.R.). Results in this report are based on work performed at beamline 23-ID-B of the Advanced Photon Source of Argonne National Laboratory, operated by UChicago Argonne, LLC, for the U.S. Department of Energy, Office of Biological and Environmental Research under contract DE-AC02-06CH11357. Computing resources from the University of Maryland Zaratan and IBBR High Performance Computing Clusters were used in this study. We thank Dr. Simon E.V. Phillips for critical reading of the manuscript.

## Author Contributions

D.W., R.Y., G.C., H.V.R.-F., M.C., and B.G.P. performed the experiments and data analyses. D.W., P.F.R., B.G.P., and R.A.M. conceived and supervised the project. All authors prepared the manuscript.

