## Supplemental Figures 1-5, Supplemental Tables 1-8 for "Structural characterization and AlphaFold modeling of human T cell receptor recognition of NRAS cancer neoantigens"

**Figures S1 to S5**

**Tables S1 to S8**

**SI References**

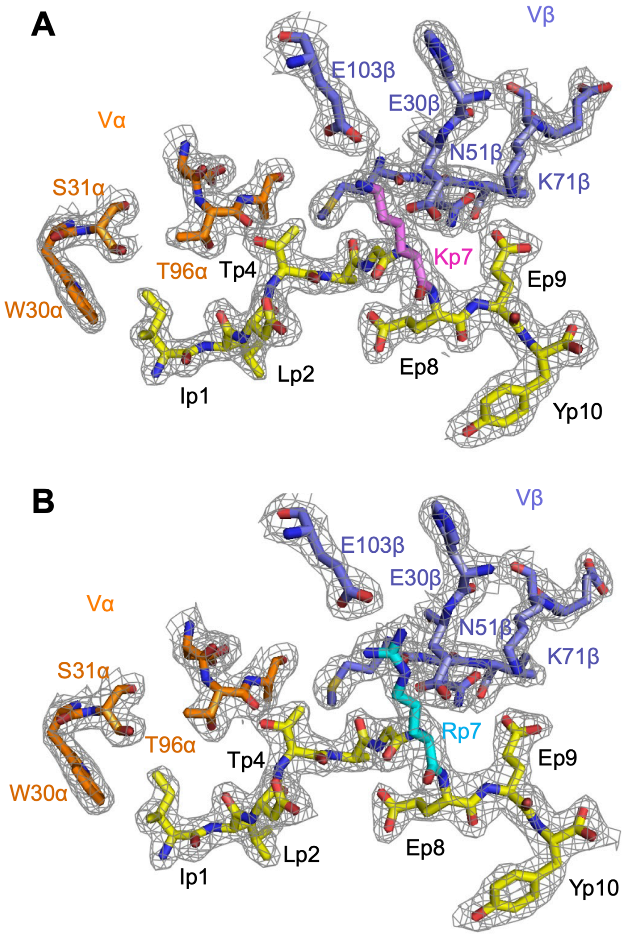

**Fig. S1.** Electron density in the interfaces of the N17.1.2–NRAS$^{Q61K}$–HLA-A1 and N17.1.2–NRAS$^{Q61R}$–HLA-A1 complexes. (*A*) Electron density in the interface of the N17.1.2–NRAS$^{Q61K}$–HLA-A1 complex. Density from the final $2F_o - F_c$ map at 2.10 Å resolution is contoured at 1σ. (*B*) Electron density in the interface of the N17.1.2–NRAS$^{Q61R}$–HLA-A1 complex. Density from the final $2F_o - F_c$ map at 2.26 Å resolution is contoured at 1σ. Carbon atoms are brown (N17.1.2 α chain), blue (N17.1.2 β chain), yellow (NRAS peptide); violet (Q61K mutant residue), or cyan (Q61R mutant residue).

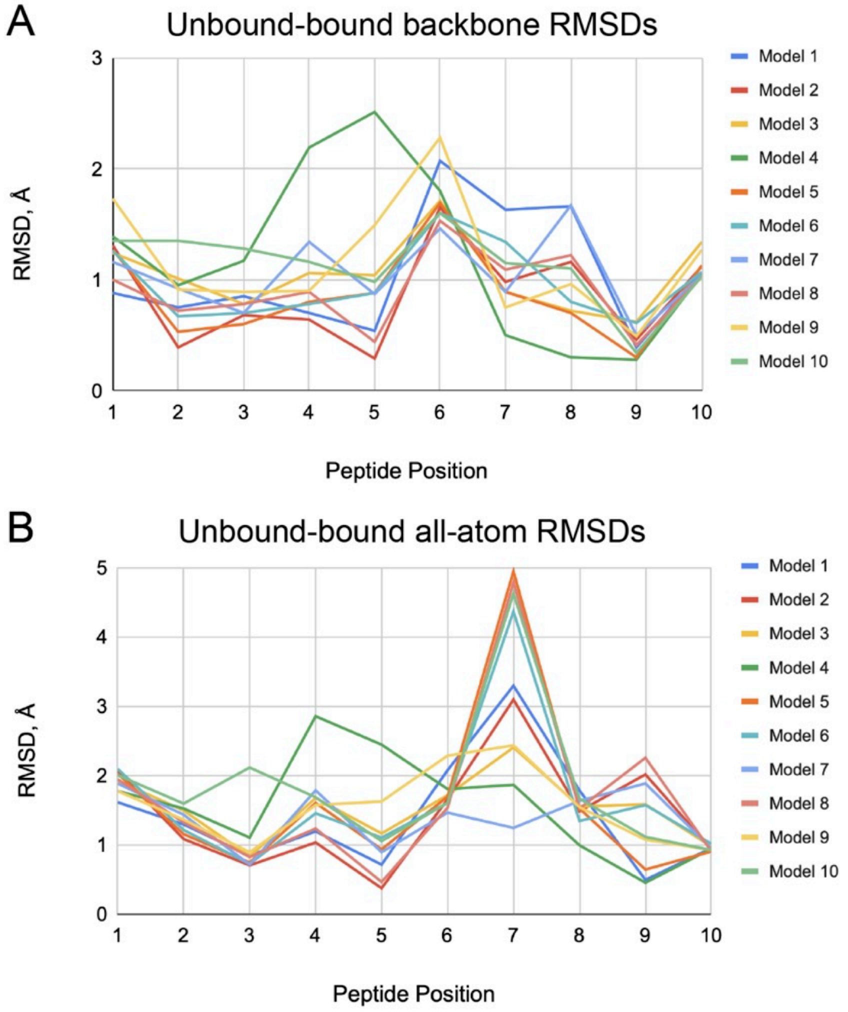

**Fig. S2.** NRAS$^{Q61K}$ peptide per-residue root-mean-square distances (RMSDs) between N17.1.2 TCR-bound structure and unbound structure NMR models. (*A*) Backbone atom and (*B*) all-atom RMSD values were calculated for each peptide position after superposition of MHC in the N17.1.2–NRAS$^{Q61K}$–HLA-A1 complex and MHC of unbound NRAS$^{Q61K}$–HLA-A1 (PDB accession code 6MPP) (1). Per-residue RMSDs were calculated between TCR-bound peptide and each of 10 NMR models of the unbound NRAS$^{Q61K}$–HLA-A1 structure (lines colored separately, as shown on right).

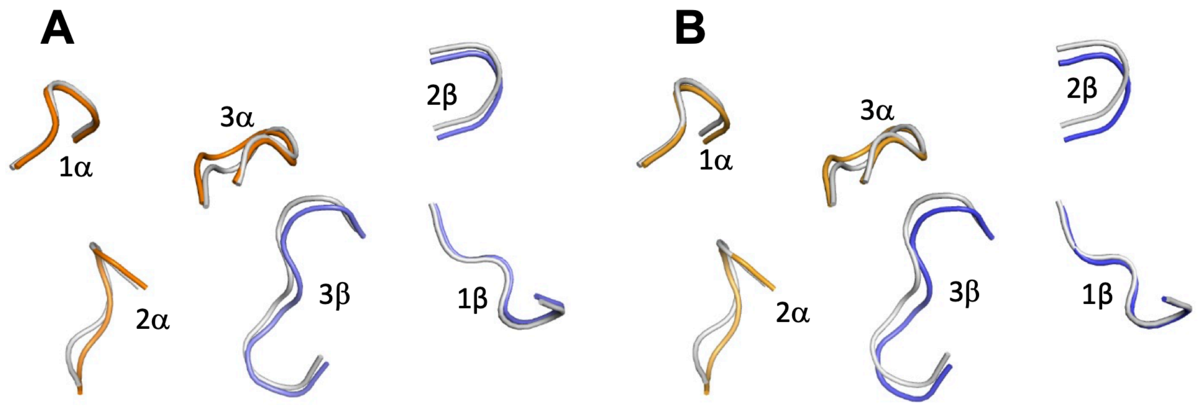

**Fig. S3.** Comparison of CDR loops in unbound versus bound TCR N17.1.2. (*A*) Superposition of CDR loops of unbound N17.1.2 (gray) onto CDR loops of N17.1.2 bound to NRAS$^{Q61K}$–HLA-A1 (Vα CDRs, orange; Vβ CDRs, blue). (*B*) Superposition of CDR loops of unbound N17.1.2 (gray) onto CDR loops of N17.1.2 bound to NRAS$^{Q61R}$–HLA-A1 (Vα CDRs, orange; Vβ CDRs, blue).

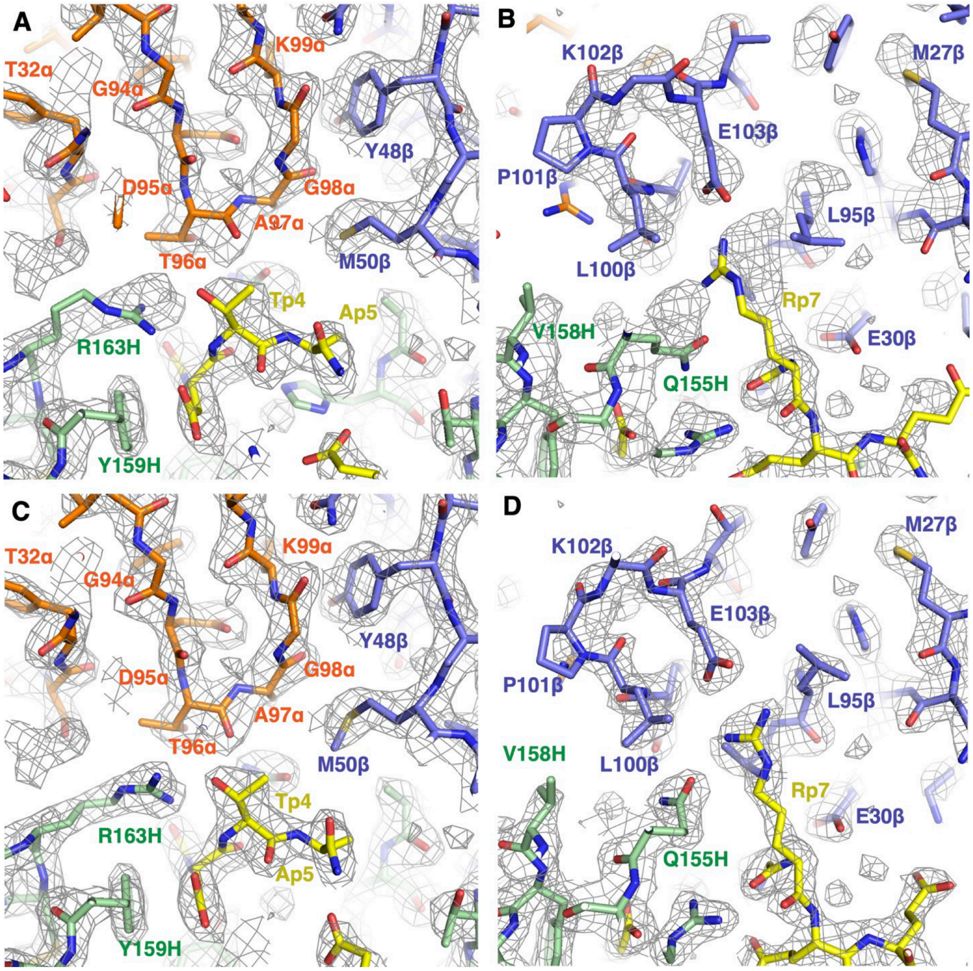

**Fig. S4.** Comparison of AlphaFold prediction of TCR N17.1.2–NRAS$^{Q61R}$–HLA-A1 complex generated by TCRmodel2 with crystallographic density map and structures. (*A*) AlphaFold prediction of N17.1.2–NRAS$^{Q61R}$–HLA-A1 complex compared with experimental electron density map in the region of CDR3α (Vα, orange; Vβ, blue; NRAS$^{Q61R}$ peptide, yellow; HLA-A1, green). (*B*) AlphaFold prediction of N17.1.2–NRAS$^{Q61R}$–HLA-A1 complex compared with electron density map in the region of CDR3β. (*C*) Model of N17.1.2–NRAS$^{Q61R}$–HLA-A1 complex built into electron density map in the region of CDR3α (Vα, orange; Vβ, blue; NRAS$^{Q61R}$ peptide, yellow; HLA-A1, green). (*D*) Model of N17.1.2–NRAS$^{Q61R}$–HLA-A1 complex built into electron density map in the region of CDR3β.

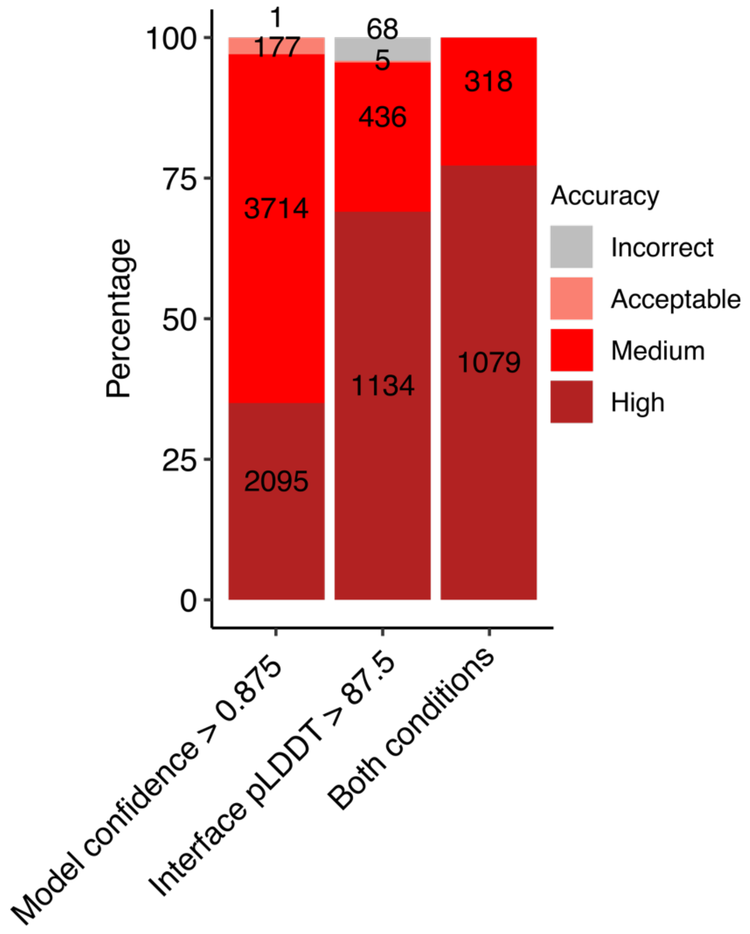

**Fig. S5.** Model accuracies when using defined AlphaFold score cutoffs for a benchmark set of TCR–pMHC complexes. For each of 20 complexes from a TCR–pMHC benchmark (2), 1000 predictions were generated and pooled for evaluation of model accuracy for the specified AlphaFold score cutoffs for model confidence, interface pLDDT (I-pLDDT), or combination of both individual cutoffs. The stacked bars illustrate the proportion of Incorrect, Acceptable, Medium, and High CAPRI accuracy criteria predictions for the cutoff. Numerical labels indicate the model counts for each accuracy level.

### Table S1. Data collection and refinement statistics

| | N17.1.2–Q61K–HLA-A1 | N17.1.2–Q61R–HLA-A1 | N17.1.2 |
|---|---|---|---|
| PDB | 8YIV | 8YJ2 | 8YJ3 |
| **Data collection** | | | |
| Resolution range (Å) | 42.7–2.10 (2.18–2.10) | 42.8–2.26 (2.34–2.26) | 47.6–3.50 (3.63–3.50) |
| Space group | *C* 2 2 21 | *C* 2 2 21 | *P* 1 21 1 |
| Unit cell parameters | 71.1, 133.2, 222.5<br>90, 90, 90 | 71.5, 134.2, 222.1<br>90, 90, 90 | 103.2, 66.1, 138.6,<br>90, 97.8, 90 |
| Total reflections[a] | 387,710 (25,546) | 464,104 (24,741) | 148,044 (14,845) |
| Unique reflections[a] | 61,763 (6,011) | 49,624 (4,524) | 23,156 (2,248) |
| Multiplicity[a] | 6.3 (4.2) | 9.3 (5.5) | 6.4 (6.6) |
| Completeness (%)[a] | 99.8 (99.0) | 98.5 (91.0) | 97.7 (97.3) |
| Mean $I/\sigma(I)$[a] | 13.7 (2.2) | 13.5 (2.2) | 9.8 (1.7) |
| Wilson *B* factor (Å$^2$) | 33.9 | 32.4 | 112.0 |
| $R_{merge}$[a,b] | 0.080 (0.403) | 0.124 (0.490) | 0.206 (0.821) |
| CC1/2 | 0.992 (0.857) | 0.985 (0.883) | 0.953 (0.777) |
| **Refinement** | | | |
| $R_{work}$[c] | 0.191 (0.228) | 0.189 (0.228) | 0.258 (0.334) |
| $R_{free}$[c] | 0.226 (0.305) | 0.229 (0.263) | 0.316 (0.366) |
| No. of protein atoms | 6,683 | 6,680 | 14,248 |
| No. of waters | 547 | 426 | 0 |
| Protein residues | 835 | 832 | 1796 |
| r.m.s.d. from ideality | | | |
| Bond lengths (Å) | 0.011 | 0.009 | 0.004 |
| Bond angles (°) | 1.46 | 1.31 | 1.10 |
| Ramachandran plot statistics | | | |
| Favored (%) | 97.0 | 97.0 | 95.0 |
| Allowed (%) | 3.0 | 3.0 | 5.0 |
| Disallowed (%) | 0 | 0 | 0 |
| Clashscore | 8.8 | 5.6 | 6.8 |
| Average *B* factor (Å$^2$) | 39.4 | 35.8 | 125.0 |
| Protein | 39.1 | 35.7 | 125.0 |
| Waters | 41.7 | 37.8 | – |

[a]Values in parentheses correspond to the highest resolution shell.

[b]$R_{merge} = \sum|I_j - \langle I \rangle|/\sum I_j$, where $I_j$ is the intensity of an individual reflection and $\langle I \rangle$ is the average intensity of that reflection.

[c]$R_{work}$ ($R_{free}$) = $\sum||F_o| - |F_c||/\sum|F_o|$; 5.0% of data were used for $R_{free}$.

**Table S2. MHC peptide groove axis position, and percentage of peptide contacts, of TCR in N17.1.2 complex structures and 82 reference TCR–peptide–MHC class I complex structures**

| Complex PDB[1] | TCR position, Å[2] | % Peptide Contacts[3] |
|---|---|---|
| 4JRY | -14.6 | 20 |
| 6TRO | -6.1 | 35 |
| **N17.1.2–Q61R–HLA-A1** | **-3.6** | 26 |
| 6ULR | -3.5 | 17 |
| **N17.1.2–Q61K–HLA-A1** | **-2.8** | 22 |
| 3FFC | -2.3 | 29 |
| 6AVF | -1.7 | 36 |
| 4QRP | -0.9 | 36 |
| 2NX5 | -0.7 | 38 |
| 8GVB | -0.5 | 19 |
| 1AO7 | -0.4 | 39 |
| 3HG1 | -0.2 | 42 |
| 5BRZ | 0.1 | 14 |
| 6R2L | 0.4 | 32 |
| 3UTS | 0.4 | 59 |
| 1KJ2 | 0.7 | 19 |
| 2AK4 | 1.2 | 68 |
| 5WKH | 1.2 | 32 |
| 3QDG | 1.3 | 40 |
| 3VXR | 1.5 | 42 |
| 3O4L | 1.7 | 31 |
| 3VXU | 1.9 | 24 |
| 6AVG | 2.0 | 40 |
| 5NHT | 2.7 | 46 |
| 3VXM | 2.7 | 60 |
| 7PHR | 2.8 | 24 |
| 7L1D | 2.9 | 27 |
| 3MV7 | 3.1 | 62 |
| 5XOV | 3.6 | 45 |
| 6RSY | 3.6 | 36 |
| 6BJ3 | 3.8 | 30 |
| 4MNQ | 3.9 | 45 |
| 6VMX | 4.0 | 37 |
| 1LP9 | 4.0 | 34 |
| 7RRG | 4.1 | 11 |
| 8DNT | 4.1 | 40 |
| 6RP9 | 4.2 | 56 |
| 6VM8 | 4.3 | 48 |
| 5JHD | 4.5 | 27 |
| 5ISZ | 4.6 | 36 |
| 3QDM | 4.9 | 51 |
| 7DZM | 5.2 | 46 |
| 6P64 | 5.2 | 50 |
| 5JZI | 5.2 | 34 |
| 5WLG | 5.3 | 27 |

| | | |
|---|---|---|
| 4MJI | 5.4 | 27 |
| 7N6E | 5.4 | 52 |
| 2YPL | 5.5 | 52 |
| 5EU6 | 5.8 | 40 |
| 1OGA | 5.8 | 35 |
| 6MTM | 5.8 | 25 |
| 4EUP | 5.8 | 56 |
| 7NDQ | 5.9 | 23 |
| 2BNQ | 5.9 | 56 |
| 1G6R | 6.0 | 49 |
| 3SJV | 6.2 | 29 |
| 7N1F | 6.2 | 58 |
| 7PDW | 6.3 | 48 |
| 2ESV | 6.3 | 45 |
| 6RPB | 6.4 | 40 |
| 5TJE | 6.4 | 40 |
| 7N2N | 6.9 | 36 |
| 8D5Q | 7.0 | 56 |
| 5D2N | 7.1 | 34 |
| 3GSN | 7.6 | 47 |
| 6RPA | 7.7 | 40 |
| 7OW5 | 7.8 | 26 |
| 1FO0 | 9.6 | 42 |
| 7RK7 | 9.6 | 38 |
| 6ZKZ | 9.7 | 23 |
| 6VRM | 9.8 | 41 |
| 5W1W | 10.4 | 30 |
| 1MI5 | 10.6 | 17 |
| 6L9L | 10.6 | 62 |
| 7N4K | 10.7 | 62 |
| 4QRR | 10.9 | 33 |
| 5IVX | 10.9 | 44 |
| 7NA5 | 11.9 | 47 |
| 7N1E | 12.0 | 59 |
| 4G8G | 12.2 | 29 |
| 3TFK | 12.4 | 33 |
| 5TEZ | 13.0 | 38 |
| 6VRN | 14.8 | 68 |
| 5SWS | 22.3 | 25 |

[1]Protein Data Bank (PDB) code of TCR–pMHC complex structure, or structure from this study (N17.1.2–Q61K–HLA-A1, N17.1.2–Q61R–HLA-A1; bold). Complexes from PDB are a set of 82 nonredundant TCR–pMHC class I structures.
[2]Position on MHC peptide groove axis, in Ångstroms, of TCR variable domain center projected onto MHC plane, as previously described (3), based on the reference frames in the TCR3d database (4). The axis is oriented with peptide C-terminus in the positive direction. Structures are listed in order of position.
[3]Percent of peptide contacts among total TCR atomic contacts (< 4 Å) to peptide and MHC.

### Table S3. Interactions between N17.1.2 and HLA-A1

| HLA-A1 | N17.1.2–Q61K | | N17.1.2–Q61R | |
|---|---|---|---|---|
| | Hydrogen bonds | Van der Waals contacts | Hydrogen bonds | Van der Waals contacts |
| α1 | | | | |
| E55H | W30α(Nε1) E55H(Oε2) | W30α(3) | W30α(Nε1) E55H(Oε2) | W30α(3) |
| E58H | S28α(Oγ) E58H(Oε2), S28α(Oγ) E58H(Oε1), W29α(Nε1) E58H(Oε1) | S28α(2), W29α(3) | S28α(Oγ) E58H(Oε2), S28α(Oγ) E58H(Oε1) | S28α(2), W29α(1) |
| Y59H | | W30α(3) | | W30α(3) |
| Q62H | T96α(N) Q62H(Oε1), T96α(Oγ1) Q62H(Oε1), T96α(Oγ1) Q62H(Nε2) | D95α(2), T96α(6) | T96α(N) Q62H(Oε1), T96α(Oγ1) Q62H(Oε1), T96α(Oγ1) Q62H(Nε2) | D95α(3), T96α(6), A97α(1) |
| R65H | Y48β(Oη) R65H(Nη1), D95α(Oδ2) R65H(Nη1), D95α(Oδ2) R65H(Nη2), | Y48β(3), D56β(6), D95α(3), A97α(3) | Y48β(Oη) R65H(Nη1), D95α(Oδ2) R65H(Nη1), D95α(Oδ2) R65H(Nη2), A97α(O) R65H(Nη1) | Y48β(3), D56β(6), D95α(3), A97α(2), K99α(1) |
| N66H | T96α(O) N66H(Nδ2) | M50β(1), T96α(3), A97α(3) | T96α(O) N66H(Nδ2) | M50β(1), T96α(4), A97α(3) |
| K68H | T55β(O) K68H(Nζ), D56β(Oδ1) K68H(Nζ), D56β(Oδ2) K68H(Nζ) | D56β(2), | T55β(O) K68H(Nζ), D56β(Oδ1) K68H(Nζ), D56β(Oδ2) K68H(Nζ) | D56β(2), |
| A69H | | M50β(1) | | M50β(1) |
| Q72H | | V52β(1), E53β(1), V54β(2) | | V52β(1), E53β(3), V54β(2) |
| T73H | | | | N51β(1) |
| α2 | | | | |
| E154H | K102β(Nζ) E154H(Oε1), K102β(Nζ) E154H(Oε2) | L100β(3), K102β(2) | K102β(Nζ) E154H(Oε2) | L100β(2), K102β(3) |
| Q155H | T98β(O) Q155H(Nε2), | V96β(3), T98β(1), L100β(5), | T98β(O) Q155H(Nε2), | V96β(2), T98β(1), L100β(2) |
| V158H | | | | L100β(1) |
| G162H | | D54α(2) | | |
| R163H | Y33α(Oη) R163H(Nη2) | S31α(2), D54α(2), T96α(7) | | S31α(1), D54α(1), T96α(8), T98β(1) |
| D166H | D54α(Oδ1) D166H(Oδ1), D54α(Oδ1) D166H(Oδ2) | W30α(1), D54α(5) | D54α(Oδ1) D166H(Oδ2) | W30α(2), D54α(9) |
| G167H | | W30α(2) | | W30α(2) |
| R170H | W30α(Nε1) R170H(Nε) | W30α(22) | W30α(Nε1) R170H(Nε) | W30α(23) |
| Y171H | | W30α(2) | | W30α(2) |

Contact residues were identified with the CONTACT program (5). Hydrogen bonds were calculated using a cut-off distance of 3.5 Å. The cut-off distance for van der Waals contacts was 4 Å.

**Table S4. TCR binding affinity changes from computational alanine scanning of HLA-A1 MHC residues**

| Mutant[1] | N17.1.2–NRAS$^{Q61K}$–HLA-A1 $\Delta\Delta G$[2] | N17.1.2–NRAS$^{Q61R}$–HLA-A1 $\Delta\Delta G$[2] |
|---|---|---|
| E55A | 1.1 | 1.2 |
| E58A | 1.0 | 1.2 |
| Y59A | 0.5 | 0.5 |
| Q62A | 1.1 | 1.1 |
| E63A | 0.2 | 0.1 |
| R65A | 2.4 | 2.4 |
| N66A | 0.6 | 0.1 |
| K68A | 0.9 | 1 |
| A69G | 0.6 | 0.6 |
| Q72A | 0.7 | 1.1 |
| T73A | 0 | 0 |
| V150A | 0.1 | 0 |
| H151A | 0 | 0 |
| E154A | 0.1 | 0.1 |
| Q155A | 1.6 | 0.8 |
| V158A | 0.1 | 0.3 |
| G162A | -0.1 | 0.1 |
| R163A | 2.1 | 2.0 |
| D166A | 0.3 | 0 |
| G167A | -0.2 | -0.2 |
| R170A | 1.1 | 1.5 |
| Y171A | 0.5 | 0.4 |

[1]MHC residue alanine substitutions for all MHC residues proximal to the TCR (< 5 Å) in the N17.1.2–NRAS$^{Q61K}$–HLA-A1 complex structure. Wild-type alanine residues were mutated to glycine.

[2]Alanine (or glycine, for wild-type alanine residues) substitutions were modeled using a protocol in Rosetta (6) to compute the TCR binding energy change ($\Delta\Delta G$). Values are in Rosetta Energy Units (REU), comparable to kcal/mol. Predicted hotspots ($\Delta\Delta G \geq 0.8$ REU) are highlighted in red.

**Table S5. Interactions between TCR N17.1.2 and NRAS$^{Q61K/R}$ peptide**

| Q61K/R | N17.1.2–Q61K | | N17.1.2–Q61R | |
|---|---|---|---|---|
| | Hydrogen bonds | Van der Waals contacts | Hydrogen bonds | Van der Waals contacts |
| I1p | | W30α(3), S31α(1) | | W30α(3), S31α(2) |
| T4p | T96α(O) T4p(Oγ1) | T96α(4) T98β(4) | T96α(O) T4p(Oγ1) | T96α(4) T98β(5) |
| G6p | | E30β(3), V96β(1) | | E30β(4) |
| K/R7p | E30β(Oε1) K7p(N), E30β(Oε2) K7p(N), E103β(Oε1) K7p(Nζ), E103β(Oε2) K7p(Nζ) | E30β(2), L95β(1), V96β(2), L100β(2), E103β(2) | E30β(Oε2) R7p(N), E103β(Oε1) R7p(Nη1), E103β(Oε2) R7p(Nη1), E103β(Oε1) R7p(Nη2) | E30β(3), L95β(2), V96β(5), L100β(2), E103β(5) |
| E9p | N51β(Oδ1) E9p(Oε2), N51β(Nδ2) E9p(Oε2), K71β(Nζ) E9p(Oε2) | N51β(1), K71β(3) | N51β(Nδ2) E9p(Oε1), K71β(Nζ) E9p(Oε1) | N51β(2), K71β(2) |

Contact residues were identified with CONTACT (5). Hydrogen bonds were calculated using a cut-off distance of 3.5 Å. The cut-off distance for van der Waals contacts was 4 Å.

**Table S6. TCR binding affinity changes from computational alanine and reversion (K61Q/R61Q) mutagenesis of peptide**

| Mutant | N17.1.2–NRAS$^{Q61K}$–HLA-A1[1] | N17.1.2–NRAS$^{Q61R}$–HLA-A1[1] |
|---|---|---|
| I55A | 0.8 | 0.7 |
| L56A | 0 | 0 |
| D57A | 0.2 | 0.3 |
| T58A | 2.1 | 2.0 |
| A59G | 0.1 | 0.1 |
| G60A | -0.2 | -0.3 |
| K61A/R61A | 1.4 | 1.8 |
| E62A | 0.2 | 0.3 |
| E63A | 0.9 | 0.9 |
| Y64A | 0 | 0 |
| K61Q/R61Q | 1.2 | 1.7 |

[1]Alanine (or glycine, for wild-type alanine residues) substitutions, or wild-type glutamine reversion substitutions (K61Q, R61Q), were modeled using a protocol in Rosetta (6) to compute the TCR binding energy change ($\Delta\Delta G$). Values are in Rosetta Energy Units (REU), comparable to kcal/mol. Predicted hotspots ($\Delta\Delta G \geq 0.8$ REU) are highlighted in red.

**Table S7. TCR–peptide polar and charged atomic contacts in X-ray and AlphaFold (TCRmodel2) modeled interfaces**

**N17.1.2–NRAS$^{Q61K}$–HLA-A1**

| Peptide residue | X-ray | TCRmodel2 |
|---|---|---|
| T4p | **T96α(O) T4p(Oγ1)** | **T96α(O) T4p(Oγ1)** |
| K7p | E30β(Oε1) K7p(N) | |
| | **E30β(Oε2) K7p(N)** | **E30β(Oε2) K7p(N)** |
| | **E103β(Oε1) K7p(Nζ)** | **E103β(Oε1) K7p(Nζ)** |
| | E103β(Oε2) K7p(Nζ) | |
| E9p | N51β(Oδ1) E9p(Oε2) | |
| | N51β(Nδ2) E9p(Oε2) | |
| | **K71β(Nζ) E9p(Oε2)** | **K71β(Nζ) E9p(Oε2)** |
| | | K71β(Nζ) E9p(Oε1) |
| | | N28β(Nζ) E9p(Oε1) |

**N17.1.2–NRAS$^{Q61R}$–HLA-A1**

| Peptide residue | X-ray | TCRmodel2 |
|---|---|---|
| T4p | **T96α(O) T4p(Oγ1)** | **T96α(O) p4T(Oγ1)** |
| R7p | **E30β(Oε2) R7p(N)** | **E30β(Oε2) R7p(N)** |
| | | E30β(Oε1) R7p(N) |
| | E103β(Oε1) R7p(Nη1) | |
| | E103β(Oε2) R7p(Nη1) | |
| | **E103β(Oε1) R7p(Nη2)** | **E103β(Oε1) R7p(Nη2)** |
| | | E103β(Oε2) R7p(Nη2) |
| E9p | N51β(Nδ2) E9p(Oε1) | |
| | **K71β(Nζ) E9p(Oε1)** | **K71β(Nζ) E9p(Oε1)** |
| | | K71β(Nζ) E9p(Oε2) |

Contacts were identified using the CONTACT program (5). Matching contacts between modeled and X-ray structures are shown on same line and in bold.

**Table S8. Confidence scores of modeled TCR–NRAS$^{Q61K}$–HLA-A1 complexes**

| | **TCRmodel2** | | **AlphaFold2.3** | | **AlphaFold3** | |
|---|---|---|---|---|---|---|
| **# Preds[1]** | **1000** | | **200** | | **5** | |
| **Complex[2]** | **Model conf[3]** | **I-pLDDT[3]** | **Model conf[3]** | **I-pLDDT[3]** | **Model conf[3]** | **I-pLDDT[3]** |
| N17.1.2 | 0.917 | 93.5 | 0.887 | 81.87 | 0.94 | 89.05 |
| N17.2 | 0.866 | 85.29 | 0.832 | 76.03 | 0.74 | 67.62 |
| N17.3.2 | 0.885 | 82.88 | 0.856 | 79.59 | 0.83 | 71.49 |
| N17.5 | 0.833 | 74.77 | 0.863 | 82.56 | 0.88 | 77.52 |
| N135.1 | 0.863 | 83.11 | 0.815 | 77.68 | 0.87 | 75.77 |

[1]Number of models generated per complex for the protocol.
[2]Complex being modeled, denoted by TCR name. All TCRs modeled in complex with NRAS$^{Q61K}$–HLA-A1 target. TCR N17.1.2 scores shown for comparison, while predictive modeling was performed for other NRAS$^{Q61K}$–HLA-A1 TCRs described in Peri et al. (7).
[3]AlphaFold model confidence and interface pLDDT (I-pLDDT) scores are shown for the top-ranked model generated for the complex with the given protocol. For AlphaFold3, the provided "ranking_score", which has a slight modification of the AlphaFold2.3 model confidence score calculation, is shown for model confidence. Cells highlighted in green in cases with both score criteria were met (model confidence > 0.875, I-pLDDT > 87.5).
